# Natural history museums are missing an opportunity to present research and collections on YouTube

**DOI:** 10.1101/2025.09.16.676522

**Authors:** Selina A. Ruzi, Adrian A. Smith, Nicole M. Lee, Nicolás J. Galvez, Timothy A. Dinh

## Abstract

There have been repeated calls in the US for scientists to communicate the importance of biological collections to both a general public audience and to policy makers. Natural history museums that house both collections and research staff while maintaining online media presences, are well positioned to communicate that value. Our study aims to understand how these museums present themselves and their research and collections via YouTube. We created a standardized and repeatable 15-question codebook to categorize content and production choices in museum account YouTube posts. In total, we analyzed 437 videos posted by 28 natural history museums in the US in 2019 and 2023, showing that messages featuring museum science are uncommon. Instead, YouTube users are likely to encounter live event recordings or promotional content related to museum or exhibit information. Research and collections themed content represents an opportunity gap for museums to engage YouTube audiences in science. If natural history museums that maintain both collections and research staff want to answer the repeated calls of communicating the importance of collections in research, then we recommend either (1) incorporating research and collections into their other video types, and/or (2) looking to other YouTube channels, some of which are subsidiary to museum institutional accounts, as a model for research and collections content strategies.

## Introduction

Natural history museums are recognized as trustworthy information centers whose missions are, generally, to serve and empower society through conserving nature (Novacek, 2008; Cameron et al., 2013). Those missions stem from a central museum function to curate collections that enable scientific research (Funk, 2018; National Academies of Sciences, Engineering, and Medicine, 2020). In recent decades, online repositories and digitization of natural history collections have revolutionized access to collections for research (Hedrick et al., 2020). However, there has simultaneously been a decline in funding and resources needed to communicate the value of museum research and collections work to decision makers and public audiences (Miller et al., 2020). Public-facing natural history museums, which have a mission of public engagement, are well positioned to communicate that value. However, little is known about how museum research and collections are represented in the public messaging of these institutions online.

The proliferation of social media publishing has presented many new opportunities for scientific institutions and individuals to communicate with public audiences (Brossard, 2013; Sugimoto et al., 2017; Jarreau et al., 2019a). In response, institutions and individuals have often pursued varying communication objectives through very different content strategies. For instance, on Instagram, Jarreau et al. (2019a) studied the #scientistswhoselfie trend and found that researchers who were posting science-themed selfies were positively impacting perceptions of trustworthiness and warmth, as well as perceptions of warmth and competence of scientists in general. In contrast, Jarreau et al. (2019b) quantitatively analyzed science museums’ postings on Instagram and “found a consistent lack of robust scientific material, science-related hashtags and scientists’ faces, among other elements that might help communicate science or encourage public participation”. Though science museums, according to Jarreau et al. (2019b), seemed to have missed an opportunity for public engagement on Instagram, communication strategies may differ on other platforms. For example, video-based platforms and users of TikTok and YouTube favor content that is produced and presented with a narrative approach and personalized perspective (Morcillo et al., 2016; Putortì et al., 2020; Huber et al., 2022). YouTube is the most well-established video-based social media platform with billions of monthly users, and over 500 hours of video content uploaded every minute (YouTube, 2024). Users searching YouTube science videos will encounter videos published by amateur enthusiasts, professional scientists, professional science communicators, and scientific institutions (Welbourne and Grant, 2016; Allgaier, 2018; Morcillo et al., 2019). To users unfamiliar with more niche YouTube science creators, institutional accounts, including those of natural history museums, are the most recognizable science-branded sources and are likely to provide primary impressions of the values of those institutions and science at-large. Therefore, the goal of this study is to understand, through a quantitative content analysis, how natural history museums are communicating on YouTube, and whether they are communicating their role in scientific research and showcasing their collections.

### Context

#### Natural History Museums and Scientific Research

The concept of what a science museum is has evolved and diversified from private collections, to academic and industrial exhibitions, and to science centers with missions of public engagement and education (Friedman, 2010; Schiele, 2021). However, specimen-based collections and the basic scientific inquiries they facilitate have been a foundational and definitive feature of museums since their inception. Today museum biological collections are the cornerstone of the scientific fields of taxonomy and systematics (Funk, 2018). Taxonomists, who discover and describe new species, often serve as curators of these museum collections. These data-rich museum specimens, beyond their involvement in taxonomic work, are used by scientists doing biodiversity research to better understand topics such as the impacts of global change (Woodward, 1987), urbanization (Shultz et al., 2020), and disease spread (Gropp, 2019).

Though the importance and value of museum collections and museum-based research is recognized among academics and museum professionals, this is not the case amongst other audiences in the US. A recent consensus report on biological collections from the National Academies of Sciences, Engineering, and Medicine (National Academies of Sciences, Engineering, and Medicine, 2020, p. 35) found that a “consistent challenge facing biological collections is a lack of awareness of the value of these collections… Ultimately, the collections community needs to improve its ability to communicate the importance of specimens in research and education to a wider audience, especially to funders and decision makers”. Even within academic institutions, the cost of maintaining these collections is often at odds with their perceived value leading to divestment in collections and closing of these museums (Davis, 2024).

Most of the largest natural history museum collections in the US are at institutions with exhibit spaces and missions that include informal education and public engagement. At those institutions, the physically exhibited collections often represent a small portion of the entire collection a museum maintains—the remainder of which are not easily accessible, housed off-exhibit and even off-site. However, digital media and the online presence of the museum can offer an expanded exhibit space to showcase the breadth and scope of these institutions’ collections and the research they facilitate.

#### Natural History Museums and Online Media

Increased connectivity via online media has transformed access to and use of natural history collections. Concerted efforts to digitize specimens and their related information have allowed researchers around the world to access collections data faster and easier than before (Dorfman, 2017; Natural-history collections face fight for survival, 2017; Ellwood et al., 2018; Gropp, 2019; National Academies of Sciences, Engineering, and Medicine, 2020). Just as researchers have used online media to share their science with one another, online media has become a primary means through which science is shared with and sought out by public audiences (Brumfiel, 2009; Brossard and Scheufele, 2013; Mitchell et al., 2016). One flourishing intersection between public science interest and collections-based research is citizen science that involves the public in data collection, curation, and/or analysis through online portals (Spear et al., 2017; Bakker et al., 2020). However, institutional social media accounts publishing public-oriented content, in the few cases that have been studied, have not been effective at engaging public audiences with natural history museum science as they rarely present original research or collections, focusing instead on promotional content (Lazzeretti et al., 2015; Jarreau et al., 2019b).

#### Science Communication on YouTube

Several studies have found specific elements of science videos that correlate with engagement and popularity metrics on YouTube, in particular strong storytelling elements, an expression of personality, and direct connection with an audience (Morcillo et al., 2016; Huang and Grant, 2020; Beautemps and Bresges, 2021). Erviti and Stengler (2016) interviewed top YouTube science content creators and reported that a strength of YouTube content is “first-hand access to researchers and experts”. First-hand science narratives, in comparison to other video styles, have been shown to be particularly effective with public audiences. For example, when academics present online videos of TED talks, those videos garner more engagement with general audiences than those by non-academics (Sugimoto et al., 2013). Additionally, Reif et al. (2020) demonstrated that scientists are perceived as more competent than YouTube science presenters though less entertaining. Similarly, in an experimental study, Ruzi et al. (2021) found that first-person video narratives from scientists received higher viewer assessments of expertise compared to third-person science presentations, and higher assessments of both trustworthiness and expertise compared to no-presenter science presentations.

The examples above show how even a simple social media strategy of putting a scientist on-screen can personify science, helping to overcome negative public perceptions of scientists (Fiske and Dupree, 2014). Personification as a strategy matches a core ethos of social media platforms like YouTube, echoed in their early motto: “Broadcast Yourself.” Therefore, a content strategy that natural history museums may apply to share their work is using first-person language with scientists or museum presenters appearing on-screen.

#### Research Questions

The overarching aim of our study is to better understand what natural history museums communicate via YouTube. We start by categorizing YouTube content from a pre-COVID time period and, for comparison, a more recent set of videos. First, because an institution’s channel can be used to host content specifically created for YouTube (i.e., videos made for YouTube audiences), but also for hosting second-hand content (e.g., live event recordings, zoom recordings; see Supplementary Material Section 3), we asked:

RQ1: To what extent are natural history museums producing and uploading content to YouTube that could be viewed as directly addressing a YouTube audience (i.e., is content created specifically for YouTube)?

Next, from the videos that do address a YouTube audience (as opposed to a previously live audience), we were interested in how often YouTube audiences might encounter content that reflects the scientific activity within these institutions. These activities might include in-progress or published scientific research related to the museum and/or its research specimen collection (see Supplementary Material Section 3). Therefore, we asked:

RQ2a: How prominent is scientific research in natural history museum YouTube content?

RQ2b: How prominent are collections and specimen-related information in natural history museum YouTube content?

Finally, we were interested in production and presentation choices related to personification, how natural history museum content is being presented to YouTube audiences, and whether those audiences are being directly called-to-action. Therefore, we asked:

RQ3: How is natural history museum YouTube content being presented to audiences relative to narrative perspective, source, and calls-to-action?

## Results

Museum accounts varied in how many total videos they had posted (mean ± SD: 275 ± 287; median: 193; range: 4 – 1,115), their total number of video views (5,628,121 ± 16,968,817; 502,092; 88 – 84,820,868), the number of subscribers (20,436 ± 63,819; 1,550; 10 – 330,000), and the number of videos scraped (13 ± 17; 7; 0 – 59) (Supplementary Table 1.1). There was a moderate strength significant positive correlation between the number of videos posted by museum accounts in both the 2019 and 2023 datasets (Spearman correlation: S = 1661.7, rho = 0.55, *p* = 0.003).

### YouTube-Specific Content

We answered RQ1 starting with the 2019 dataset which consisted of 448 videos from 28 of the 34 museums on the scraping list. Only five of the videos were not in English. Out of the remaining 443, 79% could be considered posts made for YouTube (i.e., YouTube videos or YouTube-specific content; 350 out of 443; Figure 1A). Posted content that was not initially made for YouTube consisted of recordings of presentations to physical (62 of 93 videos) or virtual audiences (31 of 93 videos; Figure 1B). Compared to those videos that were not initially produced for YouTube, YouTube videos tended to be significantly shorter content pieces (*W* = 32,356, *p* < 0.001; Table 1), were viewed significantly more (average of 3.8 times more; *W* = 13,502, *p* = 0.01; Table 1), and received 2.2 times more likes on average (*W* = 15,323, *p* = 0.39; Table 1) though the last was not statistically significant. Museum accounts varied in how many of their posts were live recordings vs. YouTube-specific content, with most posting exclusively YouTube videos or a mix of YouTube videos and live recordings (Figure 1C; Supplementary Table 1.1). When investigating the nine museum accounts that posted in both categories, 6 out of 9 still had significant differences in terms of video duration with YouTube videos being shorter, but there were only 3 out of 9 museums that had significant differences in terms of number of views and likes when accounting for false discovery rates (Supplementary Tables 5.1-5.3; Supplementary Figures 5.1-5.3).

**Fig. 1.**
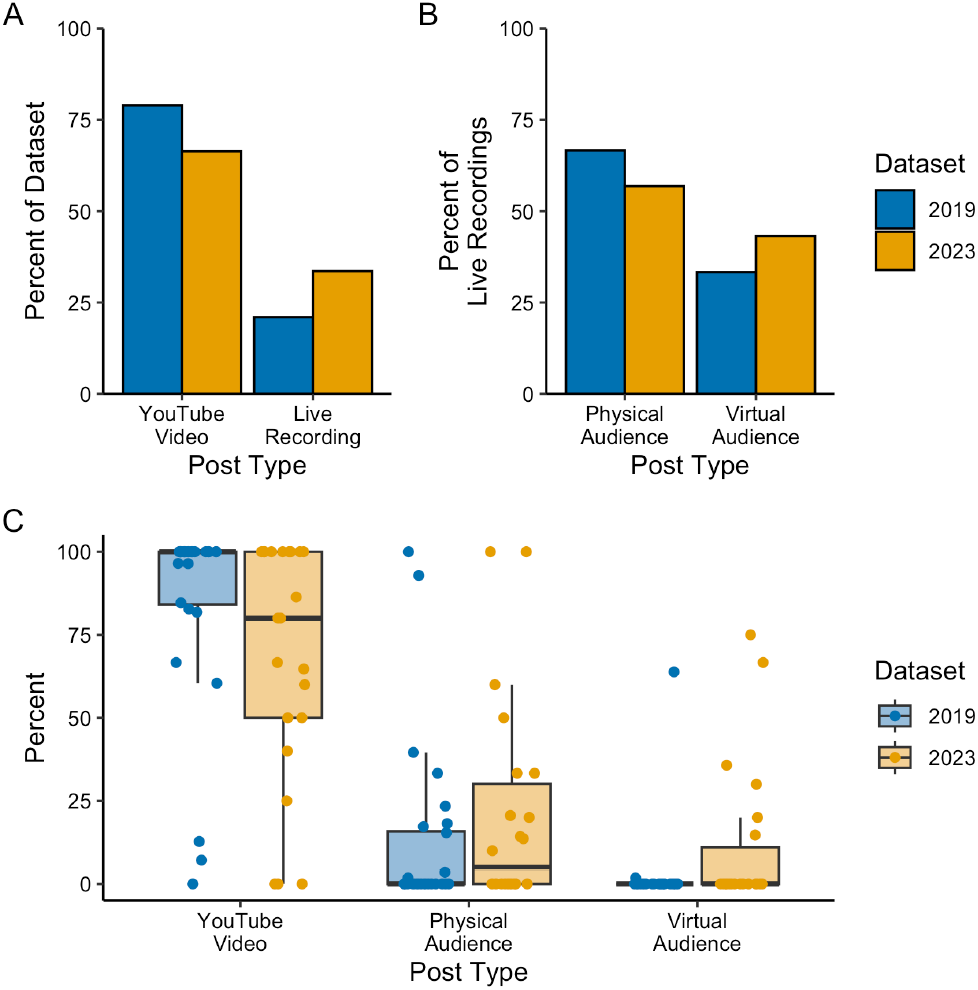
This is an example of a single column figure. < Post type by dataset. (A) Breakdown of the percent of videos that were considered either posts of YouTube videos or live recordings (sample size – videos: 2019 = 443, 2023 = 131). (B). Breakdown of the two types of live recording posts (sample size – videos: 2019 = 93, 2023 = 44). (C) Boxplots with the different percentages of the types of English videos each museum account posted in both the 2019 and 2023 datasets (sample size – museums: 2019 = 28, 2023 = 22). Points represent individual percentages from each museum jittered in space.

**Table 1.**
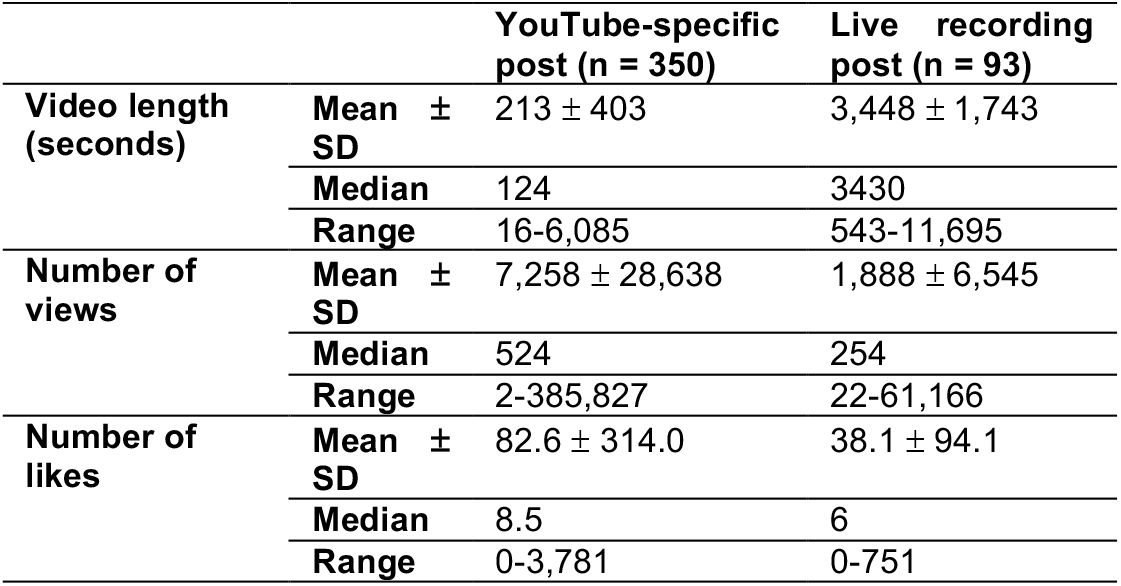
Summary of English language post type information in the 2019 dataset. The sample size, mean, standard deviation (SD), median and range of video length, number of views, and number of likes for categories of posts.

These content category trends were consistent between the 2019 and 2023 datasets (Figure 1). In the 2023 dataset, museums also varied in the percentage of their posts that were either YouTube videos or recorded live presentations to other audiences; however, these percentages were not significantly different from the 2019 dataset when accounting for false discovery rates (Figure 1C, Table 2, Supplementary Table 6.1). See the Supplementary Material Section 6 for a more detailed breakdown of the 2023 dataset.

**Table 2.** Summary of Wilcoxon tests between the 2019 and 2023 datasets. The q value accounts for false discovery rates. The degrees of freedom indicate the sample size for the 2019 and 2023 datasets respectively.

| Comparison | Statistic (W) | Degrees of freedom | p value | q value |
| --- | --- | --- | --- | --- |
| Post type: YouTube video | 214 | 28, 22 | 0.046 | 0.155 |
| Post type: Live recording – physical | 361 | 28, 22 | 0.253 | 0.479 |
| Post type: Live recording – virtual | 372 | 28, 22 | 0.052 | 0.155 |
| Main topic: sharing original research | 214 | 27, 19 | 0.266 | 0.479 |
| Main topic: sharing museum or exhibit information | 212 | 27, 19 | 0.319 | 0.479 |
| Main topic: sharing specimen or collection information | 180 | 27, 19 | 0.023 | 0.155 |
| Main topic: sharing general education | 279 | 27, 19 | 0.610 | 0.610 |
| Off-screen audio narration: no | 170 | 23, 17 | 0.488 | 0.549 |
| Off-screen audio narration: yes | 220 | 23, 17 | 0.488 | 0.549 |

### Museum Posts and Research and Collections Topics

In assessing how research (RQ2a) and collections (RQ2b) were being featured in natural history museum YouTube content, we found that four main topics could categorize museum-posted videos with our final codebook: sharing (1) original scientific research, (2) museum or exhibit information, (3) specimen or collection information, and (4) general education (Supplementary Material Section 3). Most often, museums posted to share museum or exhibit information (62.6%, or 219 out of 350). The second most common topic category was sharing general education at 17.1% (60 posts), followed by sharing original research (13.7%, 48 posts), and lastly, sharing specimen or collection information (6.6%, 23 posts) (Figure 2A). Museums varied in the categorical percentage of their posts with some museums only posting in a single topic category (sharing only museum or exhibit information with one museum only sharing general education; Figure 2B; Supplementary Table 1.1). Across museums, posting a video that shared museum or exhibit information was still the most common at 66.1% (± standard deviation-SD: ± 30.9%; range: 0.0-100.0%), followed by sharing general information (18.1 ± 23.2%; 0.0-100%), sharing original research (9.7 ± 16.2%; 0.0-63.2%), and lastly by sharing specimen or collection information (6.1 ± 10.0%; 0.0-33.3%). This trend was consistent with the 2023 dataset when accounting for false discovery rates (Figure 2A, B; Table 2; Supplementary Material Section 6).

**Fig. 2.**
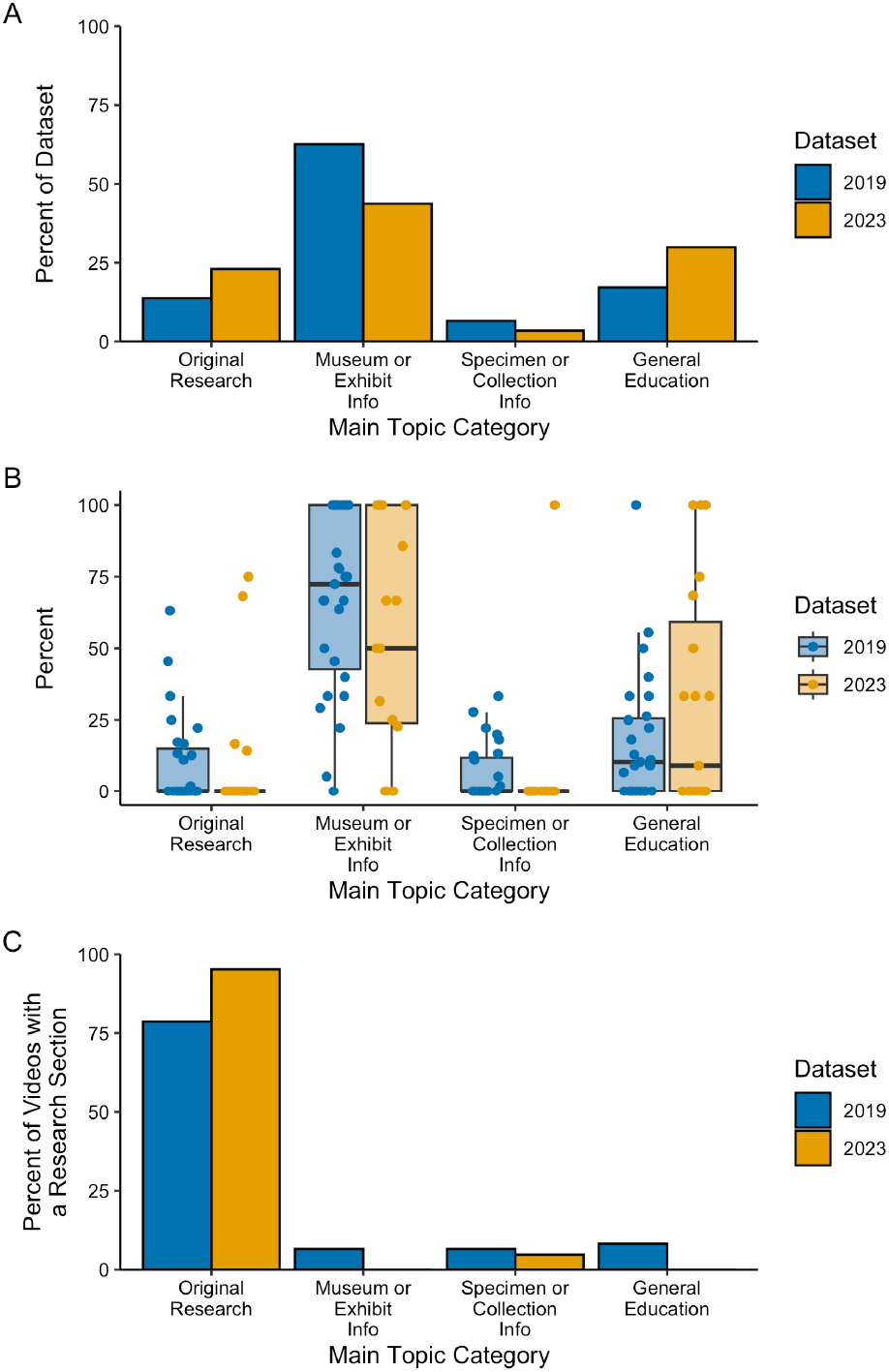
Percent of videos in each main topic category by dataset and if research is featured. (A) The overall percentage of English YouTube videos in each topic category by dataset (sample size - videos: 2019 = 350, 2023: 87). (B) Boxplots of the different percentages of English YouTube videos from each museum account that were posted in each of the main topic categories (sample size - museums: 2019 = 27, 2023 = 19). Points represent individual percentages from each museum jittered in space. (C) Percentage of YouTube videos with a research focus in each main topic category at the level of the entire dataset (sample size - videos: 2019 = 61, 2023 = 21).

There were 61 videos (17.4% of YouTube posts) that had a research focus (i.e., featured research). Based on the codebook, all videos that were categorized as a main topic category of sharing original research, by definition, also featured research. Therefore, most videos that featured research fell in the main topic category of sharing original research (48 videos; Figure 2C) though the other three main topic categories did have a few videos that also featured research. In comparison to the 2023 dataset, while there appeared to be a greater percentage of videos that featured research (24.1%, 21 videos), all but one of those videos were in the main topic category of sharing research, with the remaining one video in the main topic category of sharing specimen or collection information (Figure 2C).

Videos drew attention to collections (biological or non-biological) 46.0% (161 out of 350 videos) of the time (Figure 3A). These collections could be mentioned in any main topic category and were most often mentioned in the main topic category of sharing museum or exhibit information (95 videos) followed by sharing research (28 videos), sharing specimen or collection information (23 videos), and lastly, sharing general education (15 videos). All videos that had a main topic category of sharing specimen or collection information drew attention to collections (Figure 3B). These trends were consistent with the 2023 dataset except that sharing specimen or collection information came before sharing general education in the 2019 dataset compared to the 2023 dataset (Supplementary Material Section 6). Breaking collections down into biological and non-biological collections demonstrated that museums are more often drawing attention to biological collections than non-biological collections (Supplementary Material Section 7).

**Fig. 3.**
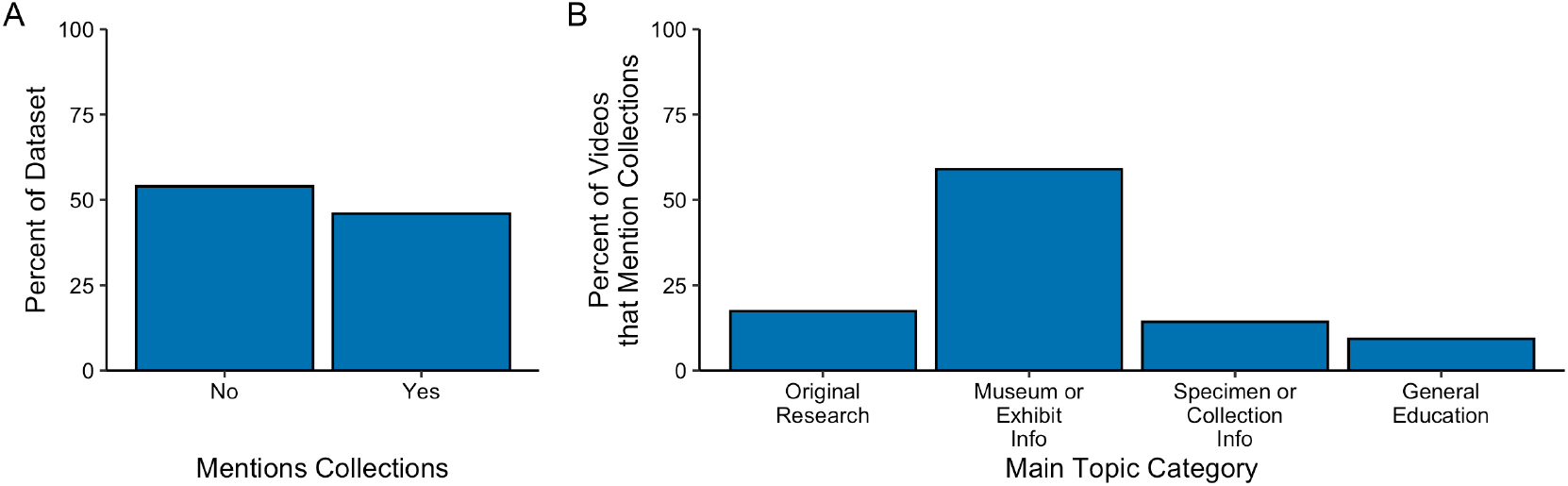
Percent of videos that draw attention to collections (biological or non-biological). (A) The overall percentage of English YouTube videos that draw attention to any collections at the level of the entire 2019 dataset (sample size – videos: 350). (B) Percentage of videos that draw attention to any collections in each main topic category at the level of the entire dataset (sample size - videos: 161)

### Presentation of Natural History Museum YouTube-Specific Content

Across all post categories and regardless of whether research was mentioned, use of at least one instance of first-person language was common (70.9%; Figure 4A). This trend was consistent within museum accounts with museums on average using at least one instance of first-person language 69.2% (± SD: ± 29.9%) of the time in their posts (Supplementary Table 1.1). This trend was consistent with the 2023 dataset (Supplementary Material Section 6).

**Fig. 4.**
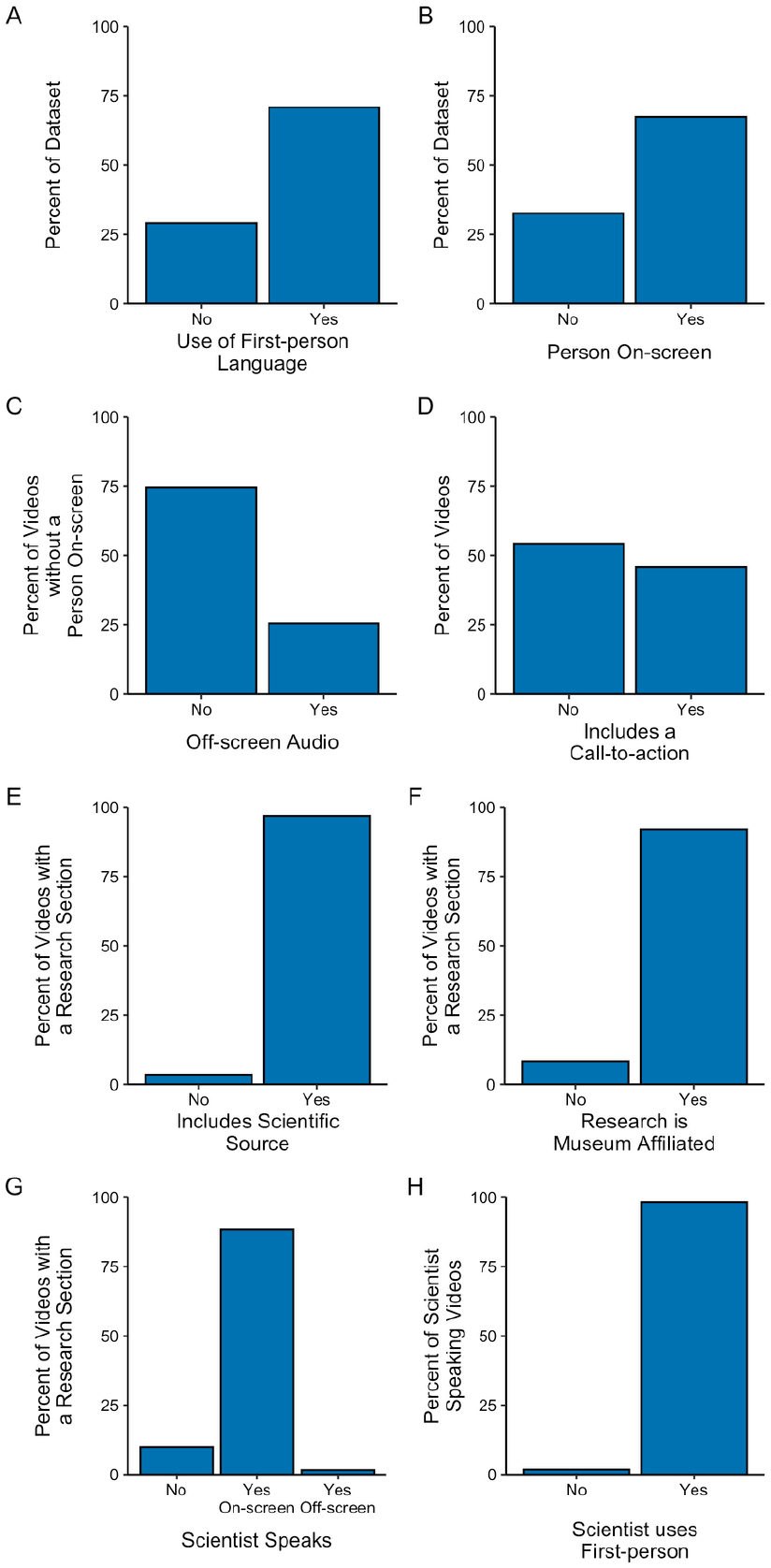
How museum YouTube content is presented. (A) The overall percentage of videos (out of 350 videos) that use at least one instance of first-person language. (B) The overall percentage of videos (out of 350 videos) that have a person speak on-screen at least some portion of the time. (C) The percentage of videos that do not have someone speak on-screen (out of 114 videos) that either do or do not have off-screen audio. (D) The percentage of videos that include a call-to-action item (out of 443 videos). (E) The percent of research videos (out of 61 videos) that include a scientific source. (F) The percent of research videos (out of 61 videos) that showcase research that is affiliated with the museum that posted the video. (G) The percent of research videos (out of 61 videos) that include a scientist speaking including if they appear on-screen or only off-screen. (H) The percent of research videos where a scientist is speaking (out of 55 videos) that use at least one instance of first-person language.

It was more common for videos to have someone speak on-screen for at least some portion of the video (67.4%) compared to having no person speak on-screen (32.6%; Figure 4B). This trend was consistent with the overall 2023 trend when looking at the dataset level. Additionally, this trend was consistent across museums (Person speaks on-screen: mean ± SD: 61.4 ± 29.1%; No person speaks on-screen: 38.6 ± 29.1%; Supplementary Table 1.1). Only two museums did not have any videos with a person speaking on-screen (Supplementary Table 1.1). For videos that did not have someone speak on-screen at any point in the video, the 2019 dataset had a higher percentage of videos that also did not have any off-screen audio narration (74.6% of videos with no one speaking on-screen) compared to the 2023 dataset which was only slightly more likely to have any off-screen audio narration (54.8%) though these differences were not significant (Figure 4C; Table 2; Supplementary Material Section 6).

Museums included call-to-action items (e.g., “click to subscribe,” “learn,” “to donate,” etc.) in 45.7% of their posts (Figure 4D) but rarely solicited feedback (e.g., ask for thoughts, replies, questions) (6 out of 160 videos). This trend was consistent with the 2023 dataset (Supplementary Material Section 6).

Most of the videos that featured research included a scientific source (59 out of 61 research videos; Figure 4E), included research that was affiliated with the museum that posted the video (56 videos; Figure 4F), had a scientist speak whether that was on-screen (54 videos) or off-screen (1 video; Figure 4G), and had that scientist use at least one instance of first-person language (54 out of 55 videos where the scientist spoke; Figure 4H). These trends were consistent with the 2023 dataset (Supplementary Material Section 6).

## Discussion

Natural history museums preserve collections of Earth’s biodiversity, both extant and extinct. The specimens they hold represent centuries of work from scientists interested in research questions describing and organizing our understanding of the natural world (Funk, 2018; Sluys, 2021). However, this discovery science and the museum infrastructure it necessitates is perpetually challenged to prove its societal value as a publicly funded asset. For example, a theme that emerged during the December 2019 (within the timeframe of our dataset) meeting of the American Institute of Biological Sciences Council of Member Societies and Organizations: “How can all scientists better quantify and demonstrate the value of biological specimens to the public and to funding agencies?” (Chang, 2020). One way museum-based scientists demonstrate their societal value is by showing connections from basic research to other research fields with immediate societal impact (Suarez and Tsutsui, 2004). Another means of publicly communicating the value of museum research is through the communication channels of the institution itself. In particular, public-facing natural history museums that have missions of both research and public engagement are well positioned for this communication challenge. While museums have many core functions, we focused specifically on institutions that have active research and maintain collections, assessing if and how museum research and collections are portrayed in the media (YouTube videos) they produce and publish online.

Our analysis of 437 English YouTube-specific videos (2019: 350; 2023: 87) posted across 28 US natural history museum accounts revealed that videos that mainly depict or present messages about research and collections are a minority (<20%) of what these museums are posting on YouTube. We found that a YouTube user going to a museum account is likely to encounter a significant (21-34%) amount of second-hand content, both pre-COVID and in a more recent year. That is, content that isn’t being presented to them, but is instead a recording of a past event, speaking to a then-live audience. Of the videos that are made to speak directly to a YouTube user, the majority of those (44-63%) are promotional content that share museum or exhibition information. While some of this content may also feature research or collections, most of it does not: out of 219 videos whose main topic categories was promoting museum or exhibit information in 2019, only 4 also included a research focus and 95 drew attention to collections. Instead, the exhibits that are promoted tend to highlight what is known rather than how it is known (i.e., does not focus on active or recent research or the scientific process) and sometimes are highlighting material that the museum has rented (i.e., traveling exhibits). Throughout our datasets, several museums were documented exclusively posting promotional content. Other studies of both natural history museums and across a wider museum field have found similar trends.

In a case study of the Museum of Natural History of Florence, Lazzeretti et al. (2015) found sharing information about collections was only mentioned once, amongst nine stated communication objectives for the museum’s posts across six social media platforms, including YouTube. They found that museum content was being generated “mostly as an instrument for communicating and promoting the museum’s activities to potential or actual visitors”. Additionally, Jarreau et al. (2019b) focused on describing science museum posts from 108 different institutional accounts on Instagram and found that 31% of posts had no science-related content, only 11% depicted a science-related professional, and less than 1% of posts provided any information about the scientific process or discovery. This practice of using social media primarily for event listings and promotional reminders has also been documented by Fletcher and Lee (2012) in US history and art museums and Suzić et al. (2016) across nearly 300 museums in Berlin and Prague. By focusing on promotional content, museums are limiting the reach of their messages. Promotional content primarily targets individuals who can attend in person events or exhibits instead of promoting online engagement and fails to take advantage of a social media platform’s potential global audience.

A more optimistic aspect of our analysis was what we found about production choices related to personification and narrative perspective. Most YouTube posts were well personified, with someone appearing on-screen speaking in first-person language. This was even more prominent in videos that featured scientific research. Though they were a minor portion (17%) of the overall 2019 set of YouTube-specific videos, nearly all of them (>88%) shared museum-affiliated research and featured a scientist on-screen presenting in first-person. If a goal of museum-generated media is to counter stereotypical perceptions of science being done by people who are unrelatable and generally lack in trust-related public perceptions (Besley et al., 2021), these production strategies have been shown to be effective in building trust in social media (Jarreau et al., 2019a; Ruzi et al., 2021). An example of this practice comes from the Carnegie Museum of Natural History, which has recently found success in terms of followers and engagement on their institutional Instagram, TikTok, and YouTube pages via regularly featuring a curator on-screen, telling jokes and sharing mollusk facts from within their research collection (Burkholder, 2021).

On institutional natural history museum YouTube channels, public audiences, in general, are not seeing research and collections as having a central role in museums and are unlikely to be hearing messages about their value within science or at a societal level. However, there are examples where those messages are being highlighted on subsidiary channels produced by groups or individuals within an institution and of museum-related content published through collaboration with other YouTube channels. Engagement seen on some of those subsidiary channels is in stark contrast to the broader institutional channels. For example, The Field Museum of Natural History has also found success through a subsidiary YouTube channel (not the main institutional channel), *The Brain Scoop*, which focuses on research and collections within the Field Museum and beyond (Dearolph, 2014; Lane, 2022). *The Brain Scoop* amassed an audience of hundreds of thousands of subscribers while The Field Museum of Natural History’s main channel, the institution that housed it, had less than ten thousand. Another example is the *Odd Animal Specimens* channel on YouTube, Instagram, and TikTok which exclusively produces content about biological museum specimens (Holdship, 2022). On YouTube, the channel has amassed over three million subscribers by showcasing specimens largely from and on-location at the University of Michigan’s Museum of Zoology, which does not have a YouTube presence of its own. An example of museum research and collection content being produced in-house for other, non-institutional channels is *Insectarium* published on the *PBS Terra* channel by PBS Digital Studios.^1^ The series features the curator of Invertebrate Zoology at the American Museum of Natural History as the host, presenting research science stories and showcasing museum-based collections. On YouTube, the eight-video series has been viewed more than 2.5 million times. A detailed comparison of the characteristics and viewership of museum YouTube content on and off institutional main channels could reveal interesting strategic content differences.

## Limitations and Future Research

Our study focused more on content-related choices than audience-related engagement metrics. Of the 350 English YouTube-specific videos we collected engagement metrics on from our 2019 dataset, the maximum number of views for a single video was 385,827 and the average single video view count was less than 8,000. These relatively low engagement numbers meant that a content analysis correlating video characteristics to viewership was unlikely to reveal insights in engagement success on the platform. This is not to say that these videos are not successfully engaging their viewers. Engagement numbers on YouTube are dependent on a suite of characteristics beyond the content itself, such as titles, thumbnails, and how the YouTube algorithm shares content on particular channels. A content analysis that incorporates engagement numbers on YouTube videos would be better suited if the comparison included videos or channels that regularly have hundreds of thousands to millions of views, and the analyses included extra content factors. Additionally, examining audience reactions was beyond the scope of this study, but future research should further examine the effects of choices such as narrative engagement on important mediating variables such as enjoyment and interest, and outcomes such as behavioral intentions.

We also did not ask how much of the content produced by museum accounts fit each respective museum’s mission statements. It is possible that content that highlights active research and collections is lacking as it is not a part of their communication plans. Future work that goes further into individual museums and analyzes how their content reflects their mission statements would be interesting. Such work should also consider taking a qualitative approach such as a semiotic intertextual analysis (Wardhani and Hasan, 2023) or discourse analysis (Lang, 2024). These methods could build upon our quantitative analysis and would be better suited to more richly describe individual channels and connect findings to semilinguistic concepts.

## Conclusions

We found that natural history museums that have research staff and collections mostly post information about the museum itself and their exhibits, and the portrayal of active research and collections has a minor role in their YouTube content. However, when they do generate science-focused content, these museums are doing a good job of providing first-hand, personified accounts of research. If raising awareness, appreciation, and increasing support for collections-based museum research is a goal for museums, there is an opportunity gap for museums to focus on producing personified depictions of the research and collections aspects of their institutions. Museums may be able to increase the recognition of their role in research and collections by incorporating some of that information into videos that otherwise serve to promote the museum without needing to dedicate entire videos to these subjects. Alternatively, looking to other YouTube channels, some that are subsidiary to their natural history museums with more focused content choices may provide an alternative model for institutional-wide accounts and content strategies in the future.

## Methods

### Museum List and Datasets

A non-comprehensive list of 34 public natural history and science museums (including two botanical gardens), varying in size and location in the US, were chosen to include in scraping (Supplementary Table 1.1). To be included, the museum had to have research staff (Supplementary Table 1.2) and maintain biological collections. We only included museums if they had both a YouTube and Twitter (now X) account to give an indication that these museums were using a variety of social media platforms to reach public audiences. We focused on museum-branded main accounts rather than subsidiary or individual researcher accounts as museum accounts would likely be the first place YouTube users would look for museum content.

For our focal dataset, we chose to evaluate posts made before COVID-19 safety measures went into effect to mitigate confounding factors due to remote work. For example, lack of in-person interactions between individual(s) in charge of social media accounts and their research colleagues could reduce the number of posts in which first-person research narratives were included. In this pre-pandemic dataset (henceforth referred to as the 2019 dataset), only 28 out of the 34 museums had posted videos within the focal time frame of 1 February 2019 to 31 January 2020 despite all having YouTube accounts (Supplementary Table 1.1). This 2019 dataset consisted of a total of 448 videos and was scraped on 22 February 2022 using Python (version 3.7.0) and YouTube’s associated application programming interface (API).

We used the list of 28 museum accounts from the 2019 dataset to generate both the training dataset and recent datasets. Both the training and recent datasets were manually compiled on 17 March 2023 and 2 August 2023, respectively. The training dataset, which is used to both help define the codebook and train the coders, consisted of up to the eight most recent videos that were published within a year of the compiling date. The recent dataset (henceforth referred to as the 2023 dataset), which was compiled for comparison to the 2019 dataset to assess whether 2019 content choices were similar or dissimilar to current trends, consisted of videos listed as being posted “3 months ago” (i.e., April 2023) up to as recently as 31 July 2023. See Supplementary Material Section 2 for a more detailed description of what information was recorded (either through scraping or manually) for each dataset.

### Codebook, Training, and Coding Procedure

Once an initial codebook was developed, we went through an iterative training process using the training dataset to ensure that coders could achieve intercoder reliability scores of 0.75 or greater for each question in the codebook (Supplementary Table 1.4). See Supplementary Material Section 3 for the training procedure.

The final codebook consisted of 15 questions that assessed various audience, content, and presentation choices (see Supplementary Material Section 3 for question wording and descriptions). Audience choices included the language the post was in (English or other), and whether the uploaded content could be viewed as being created for YouTube (YouTube video) versus second-hand content (e.g., presentation with either a physical or live audience). Content choices included what the main topic of the post was (sharing original scientific research, museum or exhibit information, specimen or collection information, or general information), whether the video had a section that featured research (yes or no), and whether attention was drawn to either biological or non-biological collections (yes or no). Presentation choices included whether first-person language was included in the media content of the post (yes or no), someone spoke on-screen (yes or no), and whether there was a specific call-to-action item (yes or no). If someone did not speak on-screen then the post was also assessed for whether there was off-screen audio narration (yes or no). If there was a call-to-action item, then the post was coded to determine if that call-to-action was to solicit comments or feedback (yes or no). Posts that were found to feature research were also assessed for the following variables: included a scientific source (yes or no), research was conducted at or was associated with the museum (yes or no), person who conducted the research was speaking in the video (yes or no), and if the person who conducted the research was in the video whether they used first-person language (yes or no).

The final codebook was then used by NJG to code both the 2019 and 2023 datasets with a 20% random subset coded by AAS to check for coder intercoder reliability (Supplementary Table 1.4). Coders were instructed to not spend more than 2 hours at a time coding to mitigate coder fatigue.

### Analyses

All analyses were conducted in R (version 4.2.1, R Core Team, 2022) and RStudio (version 2022.12.0+353 for desktop, Posit team, 2022) and graphing was conducted using the *ggplot2* package (version 3.4.0, Wickham, 2016).

Prior to conducting statistical analyses, we checked whether there was a relationship between post age and the number of views or likes (Supplementary Material Section 4). As no museum account had significant Spearman correlation between post age and number of views or likes, we did not include these variables as random effect variables in subsequent analyses (Supplementary Tables 4.1, 4.2).

We used the *wilcox_test()* function in the *rstatix* package (version 0.7.2, Kassambara, 2023) in R to run Wilcoxon sum rank tests to compare values between categories (e.g., length of video, number of views, and number of likes between YouTube videos and live recordings) and between the 2019 and 2023 datasets. These tests were sometimes done for all videos and sometimes separately for individual museum accounts. When analyses were conducted separately for individual museum accounts, we adjusted for false discovery rates by calculating *q* values (Benjamini and Hochberg, 1995).

## End Matter

## Author Contributions and Notes

Following CRediT taxonomy: SAR, AAS, NML, and NJG = conceptualization, validation; SAR, AAS, NML, NJG, and TAD = data curation, investigation, methodology, writing – review & editing; SAR and TAD = formal analysis, software, visualization; SAR, AAS, NML = funding acquisition; SAR = project administration; SAR, NML, and TAD = resources; AAS and NML supervision; SAR and AAS = writing – original draft.

The article contains supplementary information online.

All data and scripts needed to complete these analyses will be made available at reasonable request and/or when accepted at a peer-reviewed journal.

## Acknowledgments

We thank Jake Norton for assistance in collating descriptive information about the museums included in this analysis.

## Footnotes

1 https://www.pbs.org/show/insectarium/

