## Supplementary Material for "Natural history museums are missing an opportunity to present research and collections on YouTube"

#### 1 Supplemental Tables Referenced in the Main Text

**Supplementary Table 1.1. Summary information of YouTube museum accounts in the 2019 dataset.** The number of videos that were posted between 1 February 2019 and 31 January 22020 and therefore scraped, as well as a summary of the channel metrics of total number of videos posted, total number of views, and number of subscribers at time of data scraping (22 February 2022). The number of English posts in each post type. The number of videos posted and percentage of English YouTube videos (i.e., YouTube-specific content) in each of the four main topic categories, with first-person language, and with a person on-screen by museum. Num. = Number

| | | Channel metrics | | | Post type | | Main topic category | | | | | |
|---|---|---|---|---|---|---|---|---|---|---|---|---|
| **Museum (abbreviation)** | **Num. of videos scraped** | **Total num. of videos posted** | **Total num. of views** | **Num. of subscribers** | **Num. of English YouTube videos** | **Num. of English live recordings (physical presentation, virtual presentation)** | **Num. of videos sharing original research (%)** | **Num. of videos sharing museum/exhibit information (%)** | **Num. of videos sharing specimen/collection information (%)** | **Num. of videos sharing general information (%)** | **Num. of videos with any first-person language (%)** | **Num. of videos with a person on-screen (%)** |
| American Museum of Natural History (AMNH) | 33 | 1019 | 84,820,868 | 330,000 | 27 | 6 (6, 0) | 3 (11.1) | 6 (22.2) | 3 (11.1) | 15 (55.6) | 27 (100.0) | 25 (92.6) |
| Arizona Museum of Natural History (AzMNH) | 9 | 51 | 20,612 | 122 | 8 | 0 (0, 0) | 0 (0.0) | 8 (100.0) | 0 (0.0) | 0 (0.0) | 5 (62.5) | 4 (50.0) |
| Bell Museum (Bell) | 4 | 314 | 2,064,968 | 4,480 | 4 | 0 (0, 0) | 0 (0.0) | 3 (75.0) | 0 (0.0) | 1 (25.0) | 2 (50.0) | 1 (25.0) |
| Bernice P. Bishop Museum (Bishop) | 5 | 199 | 243,202 | 1,270 | 5 | 0 (0, 0) | 0 (0.0) | 2 (40.0) | 1 (20.0) | 2 (40.0) | 5 (100.0) | 4 (80.0) |

| | | Channel metrics | | | Post type | | Main topic category | | | | | |
|---|---|---|---|---|---|---|---|---|---|---|---|---|
| **Museum (abbreviation)** | **Num. of videos scraped** | **Total num. of videos posted** | **Total num. of views** | **Num. of subscribers** | **Num. of English YouTube videos** | **Num. of English live recordings (physical presentation, virtual presentation)** | **Num. of videos sharing original research (%)** | **Num. of videos sharing museum/exhibit information (%)** | **Num. of videos sharing specimen/collection information (%)** | **Num. of videos sharing general information (%)** | **Num. of videos with any first-person language (%)** | **Num. of videos with a person on-screen (%)** |
| Burke Museum of Natural History & Culture (Burke) | 15 | 213 | 1,246,962 | 4,850 | 15 | 0 (0, 0) | 2 (13.3) | 10 (66.7) | 2 (13.3) | 1 (6.7) | 9 (60.0) | 10 (66.7) |
| California Academy of Sciences (Calacademy) | 30 | 1115 | 22,594,588 | 185,000 | 24 | 5 (5, 0) | 6 (25.0) | 7 (29.2) | 3 (12.5) | 8 (33.3) | 16 (66.7) | 16 (66.7) |
| Cleveland Museum of Natural History (CMNH) | 14 | 157 | 863,027 | 3,250 | 11 | 2 (2, 0) | 5 (45.5) | 5 (45.5) | 0 (0.0) | 1 (9.1) | 10 (90.9) | 7 (63.6) |
| Delaware Museum of Nature & Science (DelMNH) | 9 | 95 | 38,628 | 151 | 9 | 0 (0, 0) | 0 (0.0) | 7 (77.8) | 2 (22.2) | 0 (0.0) | 50 (55.6) | 4 (44.4) |
| Denver Botanic Gardens (Denver Botanic) | 15 | 195 | 465,564 | 1,170 | 15 | 0 (0, 0) | 0 (0.0) | 15 (100.0) | 0 (0.0) | 0 (0.0) | 4 (26.7) | 5 (33.3) |
| Denver Museum of Nature & Science (Denver Museum) | 6 | 599 | 826,016 | 3,600 | 6 | 0 (0, 0) | 2 (33.3) | 4 (66.7) | 0 (0.0) | 0 (0.0) | 6 (100.0) | 5 (83.3) |

| | | Channel metrics | | | Post type | | Main topic category | | | | | |
|---|---|---|---|---|---|---|---|---|---|---|---|---|
| **Museum (abbreviation)** | **Num. of videos scraped** | **Total num. of videos posted** | **Total num. of views** | **Num. of subscribers** | **Num. of English YouTube videos** | **Num. of English live recordings (physical presentation, virtual presentation)** | **Num. of videos sharing original research (%)** | **Num. of videos sharing museum/exhibit information (%)** | **Num. of videos sharing specimen/collection information (%)** | **Num. of videos sharing general information (%)** | **Num. of videos with any first-person language (%)** | **Num. of videos with a person on-screen (%)** |
| Field Museum (FieldMuseum) | 0 | 271 | 1,439,499 | 6,320 | — | — | — | — | — | — | — | — |
| Florida Museum of Natural History (Florida Museum) | 18 | 226 | 885,446 | 5,000 | 18 | 0 (0, 0) | 3 (16.7) | 6 (33.3) | 5 (27.8) | 4 (22.2) | 10 (55.6) | 9 (50.0) |
| Georgia Museum of Natural History (GaMNH) | 0 | 4 | 88 | 10 | — | — | — | — | — | — | — | — |
| Harvard Museum of Natural History (Harvard) | 14 | 238 | 1,429,787 | 13,700 | 1 | 13 (13, 0) | 0 (0.0) | 1 (100.0) | 0 (0.0) | 0 (0.0) | 0 (0.0) | 0 (0.0) |
| Houston Museum of Natural History (HMNS) | 56 | 327 | 1,066,130 | 4,170 | 54 | 2 (1, 1) | 1 (1.9) | 45 (83.3) | 1 (1.9) | 7 (13.0) | 40 (74.1) | 41 (75.9) |
| Idaho Museum of Natural History (IMNH) | 6 | 23 | 939 | 20 | 4 | 2 (2, 0) | 0 (0.0) | 2 (75.0) | 0 (0.0) | 1 (25.0) | 3 (75.0) | 4 (100.0) |
| Illinois State Museum (ILState) | 4 | 114 | 64,284 | 700 | 4 | 0 (0, 0) | 0 (0.0) | 4 (100.0) | 0 (0.0) | 0 (0.0) | 4 (100.0) | 3 (75.0) |
| Kansas University Natural History | 1 | 106 | 62,043 | 309 | 1 | 0 (0, 0) | 0 (0.0) | 0 (0.0) | 0 (0.0) | 1 (100.0) | 1 (100.0) | 1 (100.0) |

| | | **Channel metrics** | | | **Post type** | | **Main topic category** | | | | | |
|---|---|---|---|---|---|---|---|---|---|---|---|---|
| **Museum (abbreviation)** | **Num. of videos scraped** | **Total num. of videos posted** | **Total num. of views** | **Num. of subscribers** | **Num. of English YouTube videos** | **Num. of English live recordings (physical presentation, virtual presentation)** | **Num. of videos sharing original research (%)** | **Num. of videos sharing museum/exhibit information (%)** | **Num. of videos sharing specimen/collection information (%)** | **Num. of videos sharing general information (%)** | **Num. of videos with any first-person language (%)** | **Num. of videos with a person on-screen (%)** |
| Museum (KUMHN) | | | | | | | | | | | | |
| La Brea Tar Pits and Museum (La Brea) | 7 | 88 | 5,271,9305 | 66,500 | 7 | 0 (0, 0) | 0 (0.0) | 7 (100.0) | 0 (0.0) | 0 (0.0) | 3 (42.9) | 2 (28.6) |
| Mace Brown Museum of Natural History (CofCNH) | 0 | 54 | 6,542 | 67 | — | — | — | — | — | — | — | — |
| Museum of the North (AlaskaMuseum) | 0 | 178 | 338,771 | 1,500 | — | — | — | — | — | — | — | — |
| Natural History Museum of Los Angeles County (NHMLA) | 59 | 818 | 3,467,896 | 8,860 | 55 | 2 (2, 0) | 7 (12.7) | 43 (78.2) | 0 (0.0) | 5 (9.1) | 40 (72.7) | 41 (74.5) |
| New Mexico Museum of Natural History & Science (NMMNHS) | 0 | 38 | 17,476 | 161 | — | — | — | — | — | — | — | — |
| New York Botanic Gardens (NYBG) | 48 | 617 | 6,175,773 | 15,800 | 29 | 19 (19, 0) | 5 (17.2) | 21 (72.4) | 0 (0.0) | 3 (10.3) | 18 (62.1) | 18 (62.1) |

| | | **Channel metrics** | | | **Post type** | | **Main topic category** | | | | | |
|---|---|---|---|---|---|---|---|---|---|---|---|---|
| **Museum (abbreviation)** | **Num. of videos scraped** | **Total num. of videos posted** | **Total num. of views** | **Num. of subscribers** | **Num. of English YouTube videos** | **Num. of English live recordings (physical presentation, virtual presentation)** | **Num. of videos sharing original research (%)** | **Num. of videos sharing museum/exhibit information (%)** | **Num. of videos sharing specimen/collection information (%)** | **Num. of videos sharing general information (%)** | **Num. of videos with any first-person language (%)** | **Num. of videos with a person on-screen (%)** |
| New York State Museum (NYSM) | 9 | 221 | 1,783,558 | 6,740 | 9 | 0 (0, 0) | 2 (22.2) | 3 (33.3) | 3 (33.3) | 1 (11.1) | 3 (33.3) | 3 (33.3) |
| North Carolina Museum of Natural Sciences (NCMNS) | 47 | 806 | 4,736,279 | 11,600 | 6 | 41 (11, 30) | 0 (0.0) | 4 (66.7) | 0 (0.0) | 2 (33.3) | 5 (83.3) | 4 (66.7) |
| Peggy Notebaert Nature Museum (Nature Museum) | 2 | 273 | 492,903 | 1,600 | 2 | 0 (0, 0) | 0 (0.0) | 1 (50.0) | 0 (0.0) | 1 (50.0) | 2 (100.0) | 2 (100.0) |
| Perot Museum of Nature & Science (Perot) | 1 | 108 | 511,280 | 1,090 | 1 | 0 (0, 0) | 0 (0.0) | 1 (100.0) | 0 (0.0) | 0 (0.0) | 0 (0.0) | 0 (0.0) |
| Sam Noble Museum (Sam Noble) | 2 | 46 | 7,471 | 84 | 2 | 0 (0, 0) | 0 (0.0) | 2 (100.0) | 0 (0.0) | 0 (0.0) | 2 (100.0) | 2 (100.0) |
| Smithsonian's National Museum of Natural History (Smithsonian) | 19 | 399 | 2,094,690 | 14,100 | 19 | 0 (0, 0) | 12 (63.2) | 1 (5.3) | 1 (5.3) | 5 (26.3) | 18 (94.7) | 17 (89.5) |
| South Carolina State Museum (SCStateMuseum) | 3 | 191 | 267,200 | 665 | 3 | 0 (0, 0) | 0 (0.0) | 3 (100.0) | 0 (0.0) | 0 (0.0) | 3 (100.0) | 1 (33.3) |

| | | Channel metrics | | | Post type | | Main topic category | | | | | |
|---|---|---|---|---|---|---|---|---|---|---|---|---|
| **Museum (abbreviation)** | **Num. of videos scraped** | **Total num. of videos posted** | **Total num. of views** | **Num. of subscribers** | **Num. of English YouTube videos** | **Num. of English live recordings (physical presentation, virtual presentation)** | **Num. of videos sharing original research (%)** | **Num. of videos sharing museum/exhibit information (%)** | **Num. of videos sharing specimen/collection information (%)** | **Num. of videos sharing general information (%)** | **Num. of videos with any first-person language (%)** | **Num. of videos with a person on-screen (%)** |
| The Academy of Natural Sciences (AcadNatSci) | 11 | 146 | 276,470 | 584 | 11 | 0 (0, 0) | 0 (0.0) | 7 (63.6) | 2 (18.2) | 2 (18.2) | 7 (63.6) | 7 (63.6) |
| Virginia Museum of Natural History (VMNH) | 0 | 37 | 17,986 | 134 | — | — | — | — | — | — | — | — |
| Yale Peabody Museum of Natural History (Peabody) | 1 | 109 | 309,866 | 1,220 | 0 | 1 (1, 0) | — | — | — | — | — | — |

**Supplementary Table 1.2. Examples of research positions at each of the museums in the datasets with rough estimates.** Examples of current research staff information including estimates (broken into three categories: <10, 10-40, 40+), some example position titles, and example programs or collections staff members may be a part of. Numbers are estimates based on current public website information (collected 2023-2024) and are intended as examples of the research performed at these museums.

| Museum | Estimate | Example Position Titles | Example Programs or Collections |
|---|---|---|---|
| American Museum of Natural History | 40+[1] | Assistant Professor, Associate Curator, Associate Director, Associate Professor, Collections Manager, Curator, Curator-in-Charge, Curatorial Assistant, Curatorial Associate, Data Collections Manager, Director, Division Chair, Field Associate, Graduate Fellow, Laboratory Manager, Museum Specialist, Ph. D. Student, Postdoctoral Fellow, Preparator, Principal Investigator, Professor, Research Associate, Resident Research Associate, Senior Museum Specialist, Senior Research Scientist, Special Collections Archivist, Special Collections and Research Librarian | African and Pacific Ethnology, Arachnida, Asian Ethnology, Astrophysics, Biological Anthropology, Center for Biodiversity & Conservation, Cnidaria, Comparative Biology, Computational Science, Crustacea, Diptera, Earth and Planetary Sciences, Fossil Mammals, Fossil Plants, Hayden Planetarium, Herpetology, Ichthyology, Institute for Comparative Genomics, Invertebrate Paleontology, Lepidoptera, Mammalogy, Minerals and Gems, Molecular Systematics, North American Ethnology, Ornithology, Paleontology, Physical Sciences, Research Library, Richard Gilder Graduate School, South American Archaeology, South American Ethnology, Vertebrate Zoology |
| Arizona Museum of Natural History | <10[2] | Collections Manager, Curator | Paleontology |

[1] https://www.amnh.org/research/staff-directory

[2] Very little information on staff lists

| Museum | Estimate | Example Position Titles | Example Programs or Collections |
|---|---|---|---|
| Bell Museum | 10-40 | Collections Manager, Curator, Science Directory & Curator | Fishes & mollusks, Genetic Resources, Herbarium, Herpetology, Mammals, Ornithology, Plants, Zoological |
| Bernice P. Bishop Museum | 10-40 | Associate Zoologist, Collections Manager, Collections Technician, Curator, Database Coordinator, Director PCMB, PCMB Research Specialist, Research Specialist, Senior Entomologist, Senior Zoologist | Botany, Entomology, Ichthyology, Invertebrate Zoology, Malacology, Marine Malacology, Molecular Biodiversity, Non-Marine Malacology, Vertebrate Zoology, Zoology |
| Burke Museum of Natural History & Culture | 40+[3] | Adjunct Curator, Affiliate Curator, Assistant Collections Manger, Collections & Lab Manager, Collections Manager, Curator, Curator Emeritus, Curatorial Associate, Field Associate, Lab Manager, Research Associate, | Archaeology, Botany, Entomology, Genetic Resources, Geology, Herpetology, Ichthyology, Malacology, Mammalogy, Micropaleontology, Ornithology, Paleobotany, Paleontology, Vertebrate Paleontology |
| California Academy of Sciences | 40+ | Assistant Curator, Biologist, Chair & Curator, Co-Director, Collections Manager, Curator, Curatorial Assistant, Department Chair & Curator, Director, Field Associate, Graduate Research, Laboratory Technician, Postdoctoral Fellow/Researcher, Programmer, Research Associate, Science Illustrator, Senior Collections Manager, Senior Curator, Senior Curatorial Assistant, Senior Laboratory Technician, Senior Research Associate, Senior Research Fellow | Anthropology, Aquatic Biology, Arachnology, Botany, Citizen Science, Comparative Genomics, Computational Biology, Entomology, Geology, Herpetology, Ichthyology, Invertebrate Zoology, Marine Biology, Mammalogy, Microbiology, Ornithology, Scientific Computing |

[3] Cultural staff not included in this number

| Museum | Estimate | Example Position Titles | Example Programs or Collections |
|---|---|---|---|
| Cleveland Museum of Natural History | 10-40 | Assistant Curator, Associate Curator, Collections Manager, Curator, Curator Emeritus, Research Associate, Senior Collections Manger, Supervisor of Field Programs | Anthropology, Archaeology, Botany, Human Health & Evolutionary Medicine, Invertebrate Paleontology, Invertebrate Zoology, Ornithology, Paleobotany and Paleontology, Vertebrate Zoology |
| Delaware Museum of Nature & Science | <10 | Assistant Curator, Collections Manager, Director of Collections & Curator, Senior Collections Manager | Birds, Mollusks |
| Denver Botanic Gardens | 10-40 | Assistant Research Scientist, Associate Curator, Associate Director, Associate Director & Curator, Associate Research Scientist, Collections Assistant, Conservation Research Associate, Curator, Director, Emeritus Curator, Head Curator, Laboratory Technician, Outreach Coordinator, Research Associate, Scientific Data Manager, Senior Curator | Alpine Plants, Applied Conservation, Aquatic Plants, Biodiversity, Ecology, Genetics, Herbology, Horticulture, Mycology, Native Plants, Natural History, Natural History Collections, Population Genetics, Seeds, Steppe Collection, Therapeutic Horticulture, Tropical Plants |
| Denver Museum of Nature & Science | 10-40 | Acting Curator, Assistant Collections Manager, Assistant Curator, Chief Preparator, Collections Assistant, Collections Manager, Curator, Digital Research Lab Technician, Director of Integrative Collections, Fossil Preparator, Head Conservator, Objects Conservator, Postdoctoral Fellow/Researcher, Preparator, Research Assistant, Research Manager, Senior Curator, Vice President | Anthropology, Archaeology, Earth Sciences, Entomology, Genetics, Geology, Health Science, Invertebrate Zoology, Ornithology, Paleobotany, Science, Space Science, Vertebrate Paleontology, Vertebrate Zoology, Vertebrates, Zoology |

| **Museum** | **Estimate** | **Example Position Titles** | **Example Programs or Collections** |
|---|---|---|---|
| Florida Museum of Natural History | 40+ | Adjunct Curator, Assistant Curator, Assistant Scientist, Associate Chair, Associate Curator, Associate Director, Associate Scientist, Biological Scientist, Collections Manager, Curator, Postdoctoral Associate, Program Director, Scientific Lab Manager | AI Biodiversity, Bioinformatics, Digital Imaging, Environmental Archaeology, Florida Archaeology, Herbarium, Herpetology, Historical Archaeology, Ichthyology, iDigBio, International Shark Observatory, Invertebrate Paleontology, Lepidoptera, Molecular Systematics, Ordway, Paleobotany, SW Florida Archaeology, Thompson Institute, Vertebrate Paleontology |
| Harvard Museum of Natural History | 40+[4] | Curator, Director, Emeritus Faculty Curator, Emeritus Honorary Curator, Research Associate | Biology, Entomology, Ichthyology, Invertebrate Paleontology, Invertebrate Zoology, Lepidoptera, Malacology, Mammalogy, Natural History, Ornithology, Vertebrate Paleontology, Zoology |
| Houston Museum of Natural History | 10-40 | Associate Curator, Consulting Curator, Curator, President & Curator | Anthropology, Astronomy, Egyptology, Entomology, Gems & Minerals, Malacology, Paleontology, Vertebrate Zoology |
| Idaho Museum of Natural History | <10[5] | Affiliate Curator, Assistant Curator, Collections Carer, Collections Manager, Curator, Directory of Marketing & Collections Manager, Director & Curator, Interim Collections Manager | Anthropology, Archaeology, Earth Sciences, Herpetology, Ichthyology, Life Science, Vertebrate Paleontology |

[4] Staff data only from Zoology branch of the museum

[5] Current and affiliated staff

| **Museum** | **Estimate** | **Example Position Titles** | **Example Programs or Collections** |
|---|---|---|---|
| Illinois State Museum | 10-40 | Assistant Curator, Collections Assistant, Curator, Registrar of Anthropology, Research Associate | Anthropology, Archaeology, Botany, Geology, Ornithology, Zoology |
| Kansas University Natural History Museum | 40+ | Adjunct Research Associate, Assistant Researcher, Associate Curator, Associate Scientist, Collection Manager, Curator, Curator Emeritus, Curator-in-Charge, Director, Doctoral Candidate, Graduate Student, Master's Student, Postdoctoral Researcher, Preparator, Research Affiliate, Researcher, Scientific Illustrator, Senior Curator, Undergraduate Researcher | Archaeology, Biodiversity, Botany, Entomology, Herpetology, Ichthyology, Informatics, Invertebrate Paleontology, Invertebrate Zoology, Mammalogy, Molecular Lab, Ornithology, Paleobotany, Phylogenetic Modeling, Vertebrate Paleontology |
| La Brea Tar Pits and Museum | 10-40 | Assistant Collections Manager, Assistant Curator, Associate Curator, Collections Manager, Curator Emeritus, Fossil Lab Manager, Preparator, Senior Collections Manager, Senior Paleontological Preparator | Paleobotanist, Paleontology |
| Natural History Museum of Los Angeles County[6] | 40+[7] | Assistant Collections Manager, Assistant Curator, Associate Collections Manager, Associate Conservator, Associate VP Collections, Collection Assistant, Collections Manager, Curator, Curator and Director, Curator Emeritus, Digitization Project Manager, Digitization Technician, Graduate Student, Postdoctoral Fellow/Researcher, Preparator, Research, Research & | Anthropology, Conservation, Crustacea, Digital Collections, Echinoderms, Entomology, Herpetology, Ichthyology, Invertebrate paleontology, Malacology, Mammalogy, Marine Biodiversity, Mineral Sciences, Ornithology, Ornithology and Mammalogy, Paleontology, Polychaetes, Vertebrate Paleontology |

[6] Some overlap with La Brea Tar Pits and Museum, tried to separate

[7] And historical curatorial staff

| Museum | Estimate | Example Position Titles | Example Programs or Collections |
|---|---|---|---|
| | | Curator, Research Assistant, Senior Collections Manager, Senior Paleontological Preparator, Senior VP Research & Collections | |
| New York Botanic Gardens[8] | 40+ | Collections Manager, Curatorial Assistant, Geographical Information Manager, Graduate Student, Herbarium Digital Asset Manager, Laboratory Manager, Laboratory Technician, Postdoctoral Research Associate, Senior Curatorial Assistant Staff Researcher | Cullman Program for Molecular Systematics, Genomics Program, Institute of Systematic Botany, William and Lynda Steere Herbarium |
| North Carolina Museum of Natural Sciences[9] | 10-40 | Assistant Head, Collections Manager, Director, Division Head, Fossil Preparator, Head, Laboratory Manager, Operations Manager, Research Adjunct, Research Curator, Senior Research Biologist | Geology, Herpetology, Ichthyology, Mammalogy, Mollusks, Non-molluscan Invertebrates, Ornithology, Paleontology |
| New York State Museum[10] | 40+ | Chief Curator, Collections Database Manager, Collections Manager, CRSP Architectural Historian, CRSP Lab Manager, CRSP Principal Investigator, Curator, Director of Research ad Collections, Education Specialist, Research and Collections Technician, Research Associate, | Anthropology, Archaeology, Bedrock Core Collection, Bioarcheology, Botany, Entomology, Ethnography, Historical Arachelogy, History, Ichthyology, Invertebrate Paleontology, Malacology, Mammalogy, Mycology, Ornithology, Paleontology, Photography Collection, |

[8] https://www.nybg.org/plant-research-and-conservation/about/science-and-conservation-staff-directory/

[9] https://naturalsciences.org/staff/rc-staff

[10] https://www.nysm.nysed.gov/research-collections/contact

| Museum | Estimate | Example Position Titles | Example Programs or Collections |
|---|---|---|---|
| | | State Archaeologist, State Entomologist | Quaternary Landscape Materials, Sedimentary Rocks, Social History, Textiles and Clothing |
| Peggy Notebaert Nature Museum[11] | <10 | Chief Curator, Curator, Senior Director of Collections, Vice President of Research and Collections | Biology, Herpetology |
| Perot Museum of Nature & Science | <10[12] | Collections Assistant, Collections Manager, Curator, Fossil Preparator, Paleo Laboratory Manager | Vertebrate Paleontology, Paleobotany |
| Sam Noble Museum | 40+ | Assistant Curator, Associate Curator, Collections Manager, Collections Manager Emeritus, Curator, Curator Emeritus, Curatorial Associate, Preparator, Research Associate | Archaeology, Herpetology, Ichthyology, Invertebrates, Invertebrate Paleontology, Mammalogy, Ornithology, Vertebrate Paleontology |
| Smithsonian's National Museum of Natural History[13] | 40+ | Adjunct Scientist, Adjunct Zoologist, Biologist, Collections Manager, Collections Assistant, Curator, Fossil Preparator, Graduate Fellow, Postdoctoral Fellow/Researcher, Research & Collections Assistant, Research Associate, Senior Researcher/Scientist | Anthropology, Botany, Entomology, Invertebrate Zoology, Mineral Sciences, Paleobiology, Smithsonian Marine Station, Vertebrate Zoology |

[11] https://naturemuseum.org/wp-content/uploads/2022/10/CAS-PNNM-Annual-Report-Fiscal-Year-2022.pdf

[12] https://www.perotmuseum.org/researchers/

[13] https://naturalhistory.si.edu/about/scientific-staff

| Museum | Estimate | Example Position Titles | Example Programs or Collections |
|---|---|---|---|
| South Carolina State Museum[14] | <10 | Collections Manager, Collections Outreach Manager, Curator | Art, Cultural History, Natural History, Pest Control, Science & Technology |
| The Academy of Natural Sciences | 40+ | Assistant Curator, Assistant Research Professor, Associate Curator, Collections Manager, Curator, Curator Emeritus, Curatorial Assistant, Doctoral Candidate, Fossil Preparator, Graduate Student, Interim Curator, Laboratory Manager, Research Associate | Botany, Diatom Herbarium, Entomology, Herpetology, Ichthyology, Invertebrate Paleontology, Malacology, Ornithology, Vertebrate Paleontology |
| Yale Peabody Museum of Natural History[15] | 40+ | Archivist, Assistant Curator, Associate Curator, Collections Manager, Curator, Curator-in-Charge, Director of Collections & Research, Museum Assistant, Museum Scientist, Senior Collection Manager, Senior Museum Assistant | Anthropology, Botany, Entomology, Herpetology, History of Science & Technology, Ichthyology, Invertebrate Paleontology, Invertebrate Zoology, Minerology & Meteoritics, Ornithology, Paleobotany, Vertebrate Paleontology, Vertebrate Zoology |

**Supplementary Table 1.3. Breakdown of the number of posts pulled for each of the three datasets.** Some posts initially pulled in the training dataset either were not used for training as the iterative rounds ended before exhausting the entire dataset and some posts were unable to be found again between pulling and coding them. The 2019 dataset (i.e., pre-pandemic dataset) includes posts from 1 February 2019 through 31 January 2020 and the 2023 dataset (i.e., recent) includes posts from April 2023 through July 2023.

[14] https://www.scstatehouse.gov/reports/aar2023/H950.pdf

[15] https://peabody.yale.edu/about/curators-collections-staff

| Museum | Training dataset | 2019 dataset | 2023 dataset |
|---|---|---|---|
| American Museum of Natural History | 8 | 33 | 3 |
| Arizona Museum of Natural History | 8 | 9 | 3 |
| Bell Museum | 8 | 4 | 1 |
| Bernice P. Bishop Museum | 7 | 5 | 1 |
| Burke Museum of Natural History & Culture | 7 | 15 | 3 |
| California Academy of Sciences | 8 | 30 | 10 |
| Cleveland Museum of Natural History | 2 | 14 | 1 |
| Delaware Museum of Nature & Science | 0 | 9 | 0 |
| Denver Botanic Gardens | 8 | 15 | 4 |
| Denver Museum of Nature & Science | 8 | 6 | 34 |
| Florida Museum of Natural History | 7 | 18 | 0 |
| Harvard Museum of Natural History | 8 | 14 | 4 |
| Houston Museum of Natural History | 8 | 56 | 5 |

| **Museum** | **Training dataset** | **2019 dataset** | **2023 dataset** |
|---|---|---|---|
| Idaho Museum of Natural History | 2 | 6 | 0 |
| Illinois State Museum | 8 | 4 | 0 |
| Kansas University Natural History Museum | 1 | 1 | 0 |
| La Brea Tar Pits and Museum | 8 | 7 | 2 |
| Natural History Museum of Los Angeles County | 8 | 59 | 24 |
| New York Botanic Gardens | 8 | 48 | 14 |
| North Carolina Museum of Natural Sciences | 8 | 47 | 5 |
| New York State Museum | 7 | 9 | 5 |
| Peggy Notebaert Nature Museum | 8 | 2 | 1 |
| Perot Museum of Nature & Science | 8 | 1 | 1 |
| Sam Noble Museum | 5 | 2 | 0 |
| Smithsonian's National Museum of Natural History | 8 | 19 | 4 |

| Museum | Training dataset | 2019 dataset | 2023 dataset |
|---|---|---|---|
| South Carolina State Museum | 5 | 3 | 1 |
| The Academy of Natural Sciences | 0 | 11 | 2 |
| Yale Peabody Museum of Natural History | 5 | 1 | 6 |

**Supplementary Table 1.4. Krippendorf's alpha coder interreliability scores for the last round of the training dataset and for the 2019 and 2023 datasets.** The last round of the training set consisted of 30 posts and was coded by four of the co-authors (SAR, AAS, NML, and NJG) while the 2019 dataset overlap consisted of 90 posts coded by two of the co-authors (NJG and AAS) and the recent dataset consisted of 27 posts coded by two of the co-authors (NJG and AAS).

| Question number | Short question description | Training dataset Krippendorf's alpha | 2019 dataset Krippendorf's alpha | 2023 dataset krippendorf's alpha |
|---|---|---|---|---|
| Q1 | Post language | 1.000 | 1.000 | 1.000 |
| Q2 | Post type | 1.000 | 0.807 | 1.000 |
| Q3 | Main topic category | 0.753 | 0.782 | 1.000 |
| Q4 | Research section | 0.908 | 0.631 | 0.882 |
| Q5 | Source for research | 0.908 | 0.729 | 0.882 |
| Q6 | Museum affiliated | 0.906 | 0.681 | 0.940 |
| Q7 | Scientist speaking in video | 0.970 | 0.761 | 0.937 |
| Q8 | Scientist uses first-person language | 0.970 | 0.736 | 0.937 |

| **Question number** | **Short question description** | **Training dataset Krippendorf's alpha** | **2019 dataset Krippendorf's alpha** | **2023 dataset krippendorf's alpha** |
|---|---|---|---|---|
| Q9 | Biological collections | 0.912 | 0.855 | 0.940 |
| Q10 | Non-biological collections | 0.835 | 0.932 | 1.000 |
| Q11 | Any first-person language | 0.904 | 0.905 | 0.942 |
| Q12 | Anyone on-screen | 0.948 | 0.945 | 1.000 |
| Q13 | Off-screen audio | 0.929 | 0.917 | 1.000 |
| Q14 | Call to action | 0.845 | 0.913 | 0.718 |
| Q15 | Solicit feedback | 0.958 | 0.968 | 0.929 |

#### 2 Datasets

A total of three different datasets were compiled for this research, the pre-pandemic dataset (henceforth referred to as the 2019 dataset), the training dataset, and the recent dataset (henceforth referred to the 2023 dataset).

##### 2.1 Pre-pandemic Dataset (Henceforth Referred to as the 2019 Dataset)

The pre-pandemic (2019) dataset consisting of 448 videos from 28 museums was scraped on 22 February 2022 using Python (version 3.7.0) and YouTube's associated application programming interface (API). The information scraped included information about the museum accounts and about individual videos. Account information included the total number of channel views, number of subscribers, total number of videos posted, account name, and account id. Video information included the video ID, title, description, video length, live or not, and if there was a transcript, what language it was in and whether the transcript was auto generated or user provided. Video engagement factor information included number of video views, number of likes, and the number of comments. Data were gathered for a total of 448 videos.

##### 2.2 Training Dataset

The training dataset was compiled manually on 17 March 2023 based on the 28 museums that had videos posted within the focal time frame of the 2019 dataset. The information that was manually recorded included the museum name, channel ID, video title, and video URL. This produced a training dataset of 176 videos, however between compiling the training dataset and training, some videos were no longer available online (at least three) and not all videos ended up being coded during training. The order of these videos was then randomized using the *sample()* function in R (version 4.2.1, R Core Team, 2022) so that during training not all the videos from one account would be coded at the same time. This dataset was used to both refine the codebook and train the coders.

#### 2.3 Recent Dataset (Henceforth Referred to as the 2023 Dataset)

The recent (2023) dataset was manually compiled on 2 August 2023 based on the 28 museums that had videos posted within the focal time frame of the 2019 dataset. This dataset was used to compare to the 2019 dataset to assess whether content choices made in the 2019 dataset were similar to current trends. Information pulled included museum name, channel ID, video title, video URL link, and verbatim post date information (i.e., "3 months ago", "5 days ago", etc.). Six museums did not have videos posted within the specified timeframe for the recent dataset.

### 3 Codebook

The "Instruction notes", "Level of analysis", and "Post variable" sections below represent the finalized codebook in both the last iteration of training and for coding both the 2019 (pre-pandemic) and 2023 (recent) datasets. The "Automatically coded variables" section describes post-processing of specific variables from the codebook.

#### 3.1 Instruction Notes

Please follow this codebook for what information to fill out and categorize for museum YouTube account posts. This file includes some examples and descriptive information to help make decisions about how to categorize posts.

Some cells may "gray" out (i.e., become gray) during the categorization process. These cells no longer need to be categorized for that post. For example, if it is determined that a post is not in English then all the remaining columns will gray out except the notes column. This is because we will not be further categorizing non-English posts. Additionally, we will be noting the presence of other YouTube video formats (e.g., presenting to a live or virtual audience) but we will not categorize past that information.

Most columns will have drop down menus with choices to choose from.

For long videos, skim through them. You can also watch them at speeds greater than normal (e.g., 2x speed).

Please use the video description to help categorize posts except for questions which specifically state to not go by the video description (e.g., do not look at the video description for questions regarding if any first-person language is included in the media content of the post, whether there is a call-to-action, or whether the post solicits feedback).

If there are questions, confusion, or comments about any aspect of this codebook please contact Selina Ruzi. Pease do not edit the "Data_validation_only" tab in the excel file.

#### 3.2 Level of Analysis

Analysis will be completed at the museum account level (e.g., AMNH, NCMNS), and at the post level (YouTube posts by different museums).

##### 3.3 Post Variables

Q1: Post language

- 1 = English (for both spoken language and if there is only text on-screen)
- 2 = Other (if other, then do not enter data into any of the other columns) (*this response will lead to all other cells graying out*)

Q2: Post type
(For Y2 and Y3, make sure to watch/skim enough of the video to double check that there is an audience. If no audience, then post type should be "1 = normal post")

- 1 = Normal post (can include video podcasts or videos that may have been recorded via zoom or another streaming platform but have been edited for content uploaded to YouTube)
- Y2 = Presentation with physical audience *(this response will lead to all other cells graying out; if speaker is in a room with an audience choose this even if screen is shared virtually)*
- Y3 = Presentation to virtual audience (i.e., zoom or other virtual meeting/stream hosting platform but should have a virtual audience component) *(this response will lead to all other cells graying out)*

Q3: Main Topic Category
(This should be on the main focus of the post so even if the post fits into more than one category, pick the one that best describes the main focus.)

- 1 = Sharing original scientific research (may include current research methodologies and expeditions)
  - Sharing the discovery of a new praying mantis species/ information on the species their scientists have described
  - Sharing work published in nature (e.g., about the face of Lucy)
  - Sharing about a restoration project that they did
  - About an expedition to collect samples (not a historic expedition)
  - Graduate students or other researchers presenting a talk about their research in conjunction with either the museum or specimens housed at the museum (while this could be thought of as a person spotlight, that it is sharing original research should be ranked more highly)
  - May use the words "New research", "study", "a study led by"
  - Could be a person talking about what they do as a researcher
  - May talk about what they study/research/work on related to science
  - *If this is chosen, then the answer to the question "Is this post about scientific research?" is "1 = Yes", the cell will highlight yellow to remind coders of this.*
  - Videos that include a lot of background, (i.e., general science education) but use this information to set up context for what they study should fall in this category (i.e., if the background leads to sharing their own research then it is "1 = sharing original scientific research")
- 2 = Museum/Exhibit Information (including exhibit information, promoting museum internships)
  - May mention "behind the scenes", "new at the museum", "visit our new exhibition all about [...]"

    - May use the words "now open" when describing a new exhibit, e.g., a promo for a new exhibit
    - Highlighting how they are updating an exhibit
    - History of the museum
    - Information about a museum program (does not matter if at museum or not)
    - Gallery view displaying multiple specimens (e.g., a tour of an exhibit)
    - Could be a video explaining how artwork for exhibitions was created; how the exhibitory art was made for display but if about a piece of art in general was made would be 3 = specimen/collection information
    - If a botanical garden creates the post, then videos of outdoor nature are considered exhibits
    - If talking about living organisms and refer to it as "our" then it is an exhibit
- 3 = Specimen/Collection information (if more about a specific specimen than an exhibit)
    - Includes both biological and non-biological specimens/collections
    - Does not include living organisms
    - May have collection or catalog numbers associated with the specimens
    - About the specific *Tyrannosaurus rex* specimen that inspired the Jurassic Park logo. Though this mentions where it is on display, it is more about the specimen itself and what this specimen inspired
    - All about a specific group of species with examples
    - About a specific specimen or how something was preserved
    - Can be about how a specific specimen was reconstructed (e.g., reconstructing an artifact)
    - Researcher talking about the history of a specific collection and highlighting its importance
    - Shows the physical specimens (may or may not be talking about them)
- 4 = General education
    - And not mainly about the specific museum, a specific researcher, or specific non-living specimen, or about sharing what a person researches or studies, etc.
    - How birds are related to dinosaurs, not about specific specimens or about sharing specific research
    - Sharing facts/information about a species (or group of species) but not a particular non-living specimen at the museum

Q4: Does the post have a section focusing on scientific research?
(either published or in progress work, need to do more than just briefly mention) (*if the main topic category was "1 = Sharing original scientific research (may include current research methodologies and expeditions, needs to be an expedition with reference to professional research science - i.e., for the purpose of scientific research and not using these words colloquially)" then the answer to this question is "1 = Yes". Cell will highlight yellow to remind of this.*

- 0 = No
- 1 = Yes

Q5: Does this research cite a specific source?
(e.g., scientists say, a specific paper or study, the person who did the research is in the video or is talking about the research, etc.) *(only answer if previous question was "1 = Yes; cell will gray out if the answer to the previous question is "0 = No")*

- 0 = No
- 1 = Yes

Q6: Is or was this scientific research conducted at the museum or by a scientist that is associated with the museum that created the post?
(i.e., scientist could have been on a field trip but works at or is affiliated with the museum, must be the museum that created the post and not a different museum. Can be affiliated with the institution, either the museum or the umbrella institution that the museum is at. Has to specifically say they are affiliated with the museum. Curator counts as affiliated with the museum. For example, if associated with any part of the Smithsonian then considered affiliated with whichever Smithsonian museum posted the content.) *(only answer if the post has a section focusing on scientific research is yes otherwise the cell will gray out)*

- 0 = No
- 1 = Yes

Q7: Is the person who did the scientific research speaking in the video?
*(only answer if the post has a section focusing on scientific research is yes otherwise the cell will gray out)*

- 0 = No
- 1 = Yes - on camera (if both on camera and voice over choose this)
- 2 = Yes - only voice over

Q8: Did the person who did the scientific research use first-person language?
*(grays out if the previous question is "0 = No")*

- 0 = No
- 1 = Yes

Q9: Does the post specifically draw attention to specimens of a biological collection? (this does not include living organisms, biological collections are dead, preserved specimens; e.g., they could be fossilized, be bones, skins, shells, specimens in amber, pinned specimens, specimens in ethanol, dried plants or fungi, seeds, and can replicas of biological specimens, etc.. If a research talk used biological specimens for the research and draws attention to them, then this should be "1 = Yes". These specimens can also be on exhibit or behind the scenes.)

- 0 = No
- 1 = Yes

Q10: Does the post specifically draw attention to specimens of a non-biological collection?
(e.g., geology – rocks, gems, minerals, etc; archeological/anthropological objects – pottery, etc.; art - paintings, sculptures, etc.)

- 0 = No
- 1 = Yes

Q11: Is any first-person language included in the media content of the post?

(YT: video only, does not include the description) (*This can include "We at the museum", any "We…" statements, "us" statement, and if the speaker introduces themselves saying "I'm…" or "My name is…"; even if this comes up only once in the media content the answer will still be "1 = Yes")*

- 0 = No
- 1 = Yes

Q12: Does at least one person speak on-screen?

- 0 = No
- 1 = Yes (*grays out next question*)

Q13: Is there any off-screen audio narration?
(i.e., is there any voice over)

- 0 = No
- 1 = Yes

Q14: Is there a call-to-action?
(e.g., does the post ask you, the viewer, to do something? Like asking people to donate, to visit, subscribe, or click a link, can include soliciting replies. Also includes sections meant to be a call and response interaction with the viewer. Do not include hashtags unless specifically called to share something using that hashtag.) (refers to the video only, does not include the video description) *(if it is text on-screen like "click to subscribe", or "learn", "to donate" counts as 1 = Yes; has to be spoken or written on-screen call to action - the speaker cannot just pause to wait for an answer if they did not tell you to "play along")*

- 0 = No
- 1 = Yes

Q15: Does the post solicit comments or feedback?
(e.g., asks for thoughts or replies or questions. The questions need to be referring to actually taking questions instead of telling you to find the answer to your questions on the website.) (refers to the video only, does not include the video description and does not matter if comments are disabled)
*(answer if Yes to previous question - if no this cell will gray out)*

- 0 = No
- 1 = Yes

Who recorded the information
*(drop down menu choice)*

Day the information was recorded
*(drop down menu choice for consistent data entry)*

Month the information was recorded
*(drop down menu choice for consistent data entry)*

Year the information was recorded
*(drop down menu choice for consistent data entry)*

Notes

- Short answer – anything coder thought may be of particular importance to note

##### 3.4 Automatically Coded Variables

As part of the final codebook, some variables would "grey" out in excel based on previous categorizations. These "greyed" out cells have meaning based on why they "grey" out. For example, for videos that were categorized in Q1 as not being in English, and in Q2 as not being a normal post type (i.e., was categorized as a live recording either to a physical or virtual audience), all other questions are automatically categorized as "NA" (Q2-Q15 or Q3-Q15 for non-English or live recordings respectively) as it is not possible to know what traits the videos would have for each of the subsequent questions. The same goes for when Q12 is categorized as having anyone speaking on-screen, as Q13 assessing whether there was off-screen audio "greys" out. As Q13 is not assessed and it is possible for someone to appear on-screen speaking only some of the time and also speak off-screen, or have another individual speak off-screen, we cannot be sure that these videos have or do not have off-screen audio narration. Therefore, these videos would have "NA" entered for Q13. However, there are also instances where categorizing one question will "grey" out other questions though we do know that the video will not have these characteristics. This is the case for videos that do not feature research (Q4). If a video does not feature research then it will not have source for the research (Q5), there will not be any museum affiliation for the research (Q6), a scientist will not be speaking in the video (Q7), and the scientist would not be using first-person language (Q8). Therefore Q5-Q8 would all have an automatic answer of "no" for when there is no research focus. Additionally, even if the post featured research (Q4), but there was no scientist speaking in the video (Q7) then the scientist would not have been using first-person language in the video (Q8 = "no"). As part of the codebook, we also consider soliciting questions or feedback (Q15) as a subset of a video including a call to action (Q14), therefore if there is no call to action (Q14 = "no"), the "greyed" out Q15 is automatically "no". We use R (version 4.2.1, R Core Team, 2022) and RStudio (version 2022.12.0+353 for desktop, Posit team, 2022) to fill in these responses in the "greyed" out cells prior to assessing intercoder reliability using either ReCal3[16] (training dataset) or ReCal2[17] (pre-pandemic and recent datasets) with "NA" set to "99" and "no" set to "0" as both ReCal3 and ReCal2 requires numerical digits for categorizations.

##### 3.5 Training

After an initial codebook was developed, we went through an iterative training process to both refine the codebook and ensure that high intercoder reliability scores could be achieved. The iterative process consisted of co-authors coding from 10 to 33 of posts from the training dataset, checking for intercoder reliability, editing the codebook to improve either questions or descriptions on how to categorize posts, and then starting the next iteration of coding based on the updated codebook. Co-authors SAR, AAS, NML, and NJG all took part in the iterative training process. In total, we went through six iterations coding a total of 113 posts before settling on the final codebook. The sixth

[16] https://dfreelon.org/utils/recalfront/recal3/

[17] http://dfreelon.org/utils/recalfront/recal2/

iteration of coding consisted of 30 posts using the final codebook and achieved intercoder reliability scores across all four coders of 0.753+ for each of the 15 codebook questions (Supplementary Table 1.4).

#### 4 Age of Posts

We assumed that because videos were scraped almost two years (on 22 February 2022) after the end of the focal period of posting (31 January 2020) that time was not a significant factor in engagement differences. To check this assumption, we conducted Spearman correlations using the *cor.test()* function in R for both number of views and number of likes. We did this at both the level of all videos scraped and within museum accounts that had posted at least 10 videos each. To account for multiple analyses when investigating correlations at the museum account level, we calculated *q* values to account for false discovery rates (Benjamini and Hochberg, 1995).

##### 4.1 Calculating the Age of Posts

We determined the age of the videos based on the number of days that had passed since the video was published on YouTube to when the data was scraped. Higher numbers indicated the video was posted closer to the beginning of the focal time frame and lower numbers indicated the video was posted closer to the end of the focal time frame.

##### 4.2 Age and Number of Views

Irrespective of museum account and whether the video was in English or considered a YouTube-specific post, there was a significant but weak Spearman correlation of video age with number of views videos received ($S$ = 12940108, rho = 0.14, $p$ value = 0.004). However, the strength of this correlation did vary by museum account that had posted at least 10 videos and some museums even had a slight negative correlation. However, after adjusting for false discovery rates, no museum account had a significant correlation between video age and number of views (Supplementary Table 4.1). Prior to accounting for false discovery rates, only two museums had a significant correlation between video age and number of views: Florida Museum and New York Botanical Garden.

**Supplementary Table 4.1. Summary of Spearman correlations of video age with number of views.** *q* values were calculated using the *p.adjust* function in R (version 4.2.1, R Core Team, 2022) using RStudio (version 2022.12.0+353 for desktop, Posit team, 2022) to account for false discovery rates.

| **Museum** | ***S*** | **rho** | ***p* value** | ***q* value** |
|---|---|---|---|---|
| American Museum of Natural History | 5432 | 0.09 | 0.61 | 0.79 |
| Burke Museum | 433 | 0.23 | 0.42 | 0.79 |
| California Academy of Sciences | 5638 | -0.25 | 0.18 | 0.46 |
| Cleveland Museum of Natural History | 652 | -0.43 | 0.12 | 0.43 |

| | | | | |
|---|---|---|---|---|
| Denver Botanic Gardens | 641 | -0.15 | 0.61 | 0.79 |
| Florida Museum | 490 | 0.50 | 0.04 | 0.38 |
| Harvard Museum of Natural History | 484 | -0.06 | 0.83 | 0.83 |
| Houston Museum of Natural Science | 31490 | -0.08 | 0.58 | 0.79 |
| Natural History Museum of Los Angeles County | 32976 | 0.04 | 0.79 | 0.83 |
| New York Botanical Garden | 13341 | 0.28 | 0.06 | 0.38 |
| North Carolina Museum of Natural Sciences | 13456 | 0.22 | 0.13 | 0.43 |
| Smithsonian's National Museum of Natural History | 1328 | -0.17 | 0.50 | 0.79 |
| The Academy of Natural Sciences | 236 | -0.07 | 0.83 | 0.83 |

##### 4.3 Age and Number of Likes

Similar to the results from the number of views, irrespective of museum account and whether the video was in English or considered a normal post, there was a significant but weak Spearman correlation of video age with the number of likes videos received ($S$ = 12856257, rho = 0.14, $p$ value = 0.003). Again, the strength of this correlation did vary by museum account that had posted at least 10 videos and some museums even had a slight negative correlation, and no correlation was significant after accounting for false discovery rates (Supplementary Table 4.2). Prior to accounting for false discovery rates, three museums had significant correlations between video age and the number of likes. This again included the New York Botanical Garden, but this time the other museums included the American Museum of Natural History and the California Academy of Sciences.

**Supplementary Table 4.2. Summary of Spearman correlations of video age with number of likes.** *q* values were calculated using the *p.adjust* function in R to account for false discovery rates.

| **Account** | ***S*** | **rho** | ***p* value** | ***q* value** |
|---|---|---|---|---|
| American Museum of Natural History | 3795 | 0.37 | 0.04 | 0.17 |
| Burke Museum | 523 | 0.07 | 0.81 | 0.91 |
| California Academy of Sciences | 6383 | -0.42 | 0.02 | 0.17 |

| | | | | |
|---|---|---|---|---|
| Cleveland Museum of Natural History | 664 | -0.46 | 0.10 | 0.25 |
| Denver Botanic Gardens | 727 | -0.30 | 0.28 | 0.52 |
| Florida Museum | 565 | 0.42 | 0.09 | 0.25 |
| Harvard Museum of Natural History | 481 | -0.06 | 0.85 | 0.91 |
| Houston Museum of Natural Science | 28809 | 0.02 | 0.91 | 0.91 |
| Natural History Museum of Los Angeles County | 30534 | 0.11 | 0.42 | 0.64 |
| New York Botanical Garden | 12910 | 0.30 | 0.04 | 0.17 |
| North Carolina Museum of Natural Sciences | 13592 | 0.21 | 0.15 | 0.32 |
| Smithsonian's National Museum of Natural History | 1355 | -0.19 | 0.44 | 0.64 |
| The Academy of Natural Sciences | 175 | 0.20 | 0.55 | 0.71 |

##### 4.4 Summary

When accounting for false discovery rates, no museum account had significant Spearman correlations between the age of the post and number of views or likes; therefore, we do not include number of views or likes as random effect variables in subsequent analyses (Supplementary Tables 4.1, 4.2).

#### 5 Engagement Factors

##### 5.1 Video Duration

Video duration was measured in seconds and was included in the data that was scraped directly from YouTube using YouTube's associated application programming interface (API) for the 2019 dataset. Live recording posts had longer video durations than YouTube-specific videos both when investigating at the entire dataset level ($W = 32,356$, $p < 0.001$) and for 6 of the 9 museum accounts that had posted both YouTube specific videos and of live recordings (Supplementary Table 5.1, Supplementary Figure 5.1). One additional museum (Cleveland Museum of Natural History) had a marginally significant result when taking into account false discovery rates (Supplementary Table 5.1). In all cases, Supplementary Figure 5.1 demonstrates that there is a trend for live recording posts to be longer than YouTube-specific videos.

**Supplementary Table 5.1. Summary of Wilcoxon tests for video duration between YouTube-specific and live recording posts by museum account.** The degrees of freedom indicate the sample size for the live recordings and YouTube videos respectively. False discovery rates take into account tests run on video duration, number of video views, and number of likes videos received. **Bold** values are significant.

| **Museum (abbreviation)** | **Statistic (*W*)** | **Degrees of freedom** | ***p* value** | ***q* value** |
|---|---|---|---|---|
| American Museum of Natural History (AMNH) | 161.0 | 6, 27 | **< 0.001** | **< 0.001** |
| California Academy of Sciences (Calacademy) | 120.0 | 5, 24 | **< 0.001** | **0.003** |
| Cleveland Museum of Natural History (CMNH) | 22.0 | 2, 11 | 0.026 | 0.053 |
| Harvard Museum of Natural History (Harvard) | 13.0 | 13, 1 | 0.143 | 0.214 |
| Houston Museum of Natural History (HMNS) | 108.0 | 2, 54 | **0.018** | **0.044** |
| Idaho Museum of Natural History (IMNH) | 8.0 | 2, 4 | 0.133 | 0.211 |
| Natural History Museum of Los Angeles County (NHMLA) | 109.0 | 2, 55 | **0.020** | **0.046** |
| New York Botanical Garden (NYBG) | 551.0 | 19, 29 | **< 0.001** | **< 0.001** |
| North Carolina Museum of Natural Sciences (NCMNS) | 246.0 | 41, 6 | **< 0.001** | **< 0.001** |

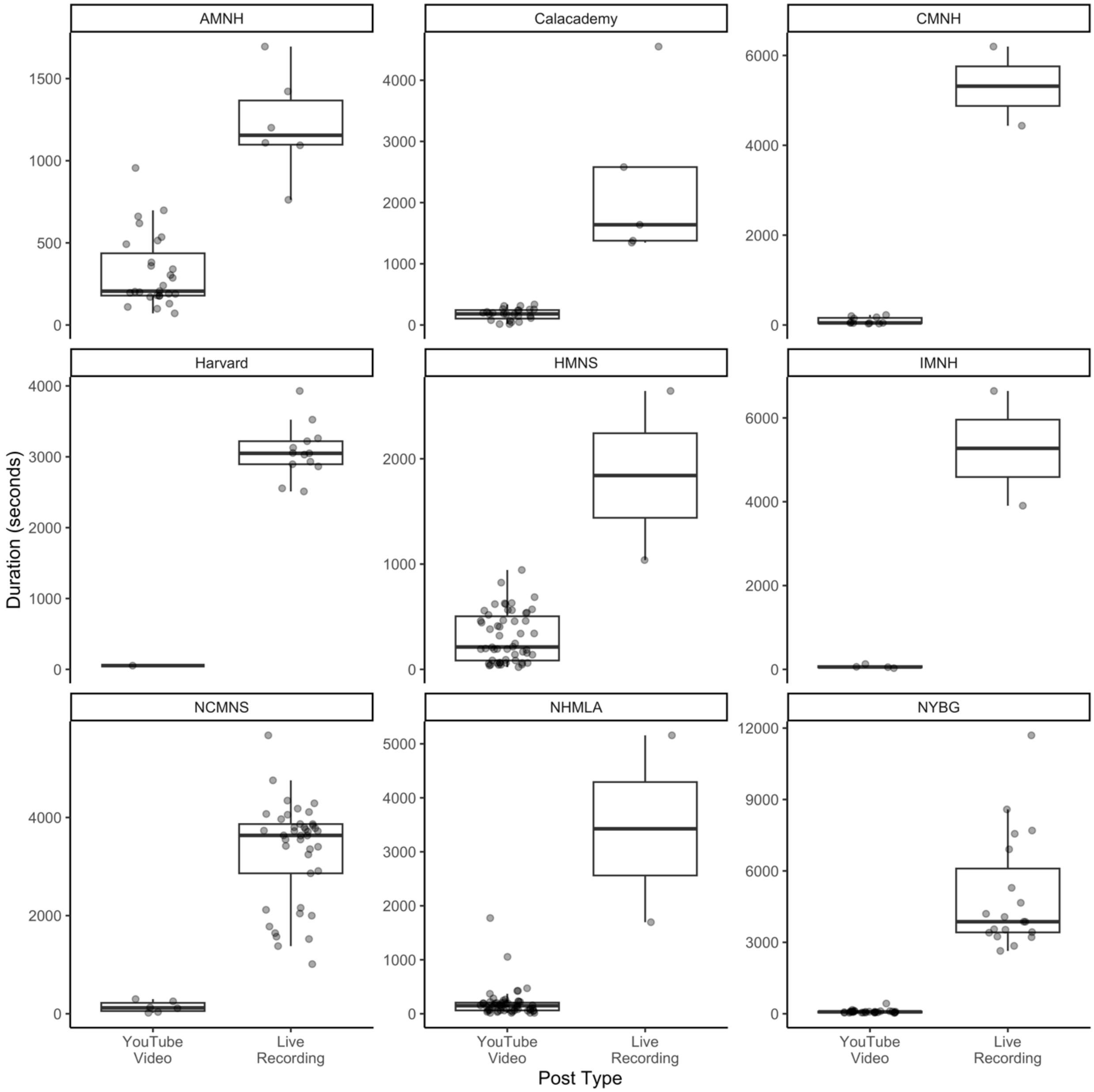


**Supplementary Figure 5.1. Boxplots of video duration (seconds) between YouTube-specific and live recording posts by museum account.** Points represent values from individual videos from each museum jittered horizontally in space. Outliers for the boxplot are suppressed. See Supplementary Table 1.1 for museum acronyms and sample sizes (i.e., number of normal and non-normal videos) for each museum account.

#### 5.2 Number of Views

Number of video views as of 22 February 2022 was scraped directly from YouTube using YouTube's associated application programming interface (API) for the 2019 dataset. Live recordings tended to have fewer views than YouTube-specific videos when investigating at the entire dataset level ($W$ = 13,502, $p$ = 0.01) but were only significantly different for 3 of the 9 museum accounts that had posted both YouTube-specific videos and live recordings (Supplementary Table 5.2,

Supplementary Figure 5.2). Interestingly, for some museums that posted both YouTube-specific videos and live recordings the live recordings had a higher number of views, though these differences were not significant. The three museums accounts that were significant all demonstrate that YouTube-specific videos had a higher view count than live recording posts.

**Supplementary Table 5.2. Summary of Wilcoxon tests for number of views between YouTube-specific and live recording posts by museum account.** The degrees of freedom indicate the sample size for the live recordings and YouTube-specific videos respectively. False discovery rates take into account tests run on video duration, number of video views, and number of likes videos received. **Bold** values are significant.

| **Museum (abbreviation)** | **Statistic (*W*)** | **Degrees of freedom** | ***p* value** | ***q* value** |
|---|---|---|---|---|
| American Museum of Natural History (AMNH) | 0.0 | 6, 27 | **< 0.001** | **< 0.001** |
| California Academy of Sciences (Calacademy) | 13.0 | 5, 24 | **0.004** | **0.014** |
| Cleveland Museum of Natural History (CMNH) | 20.0 | 2, 11 | 0.103 | 0.185 |
| Harvard Museum of Natural History (Harvard) | 7.0 | 13, 1 | 1.000 | 1.000 |
| Houston Museum of Natural History (HMNS) | 42.0 | 2, 54 | 0.612 | 0.711 |
| Idaho Museum of Natural History (IMNH) | 8.0 | 2, 4 | 0.133 | 0.211 |
| Natural History Museum of Los Angeles County (NHMLA) | 80.0 | 2, 55 | 0.288 | 0.389 |
| New York Botanical Garden (NYBG) | 115.5 | 19, 29 | **< 0.001** | **0.003** |
| North Carolina Museum of Natural Sciences (NCMNS) | 107.5 | 41, 6 | 0.632 | 0.711 |

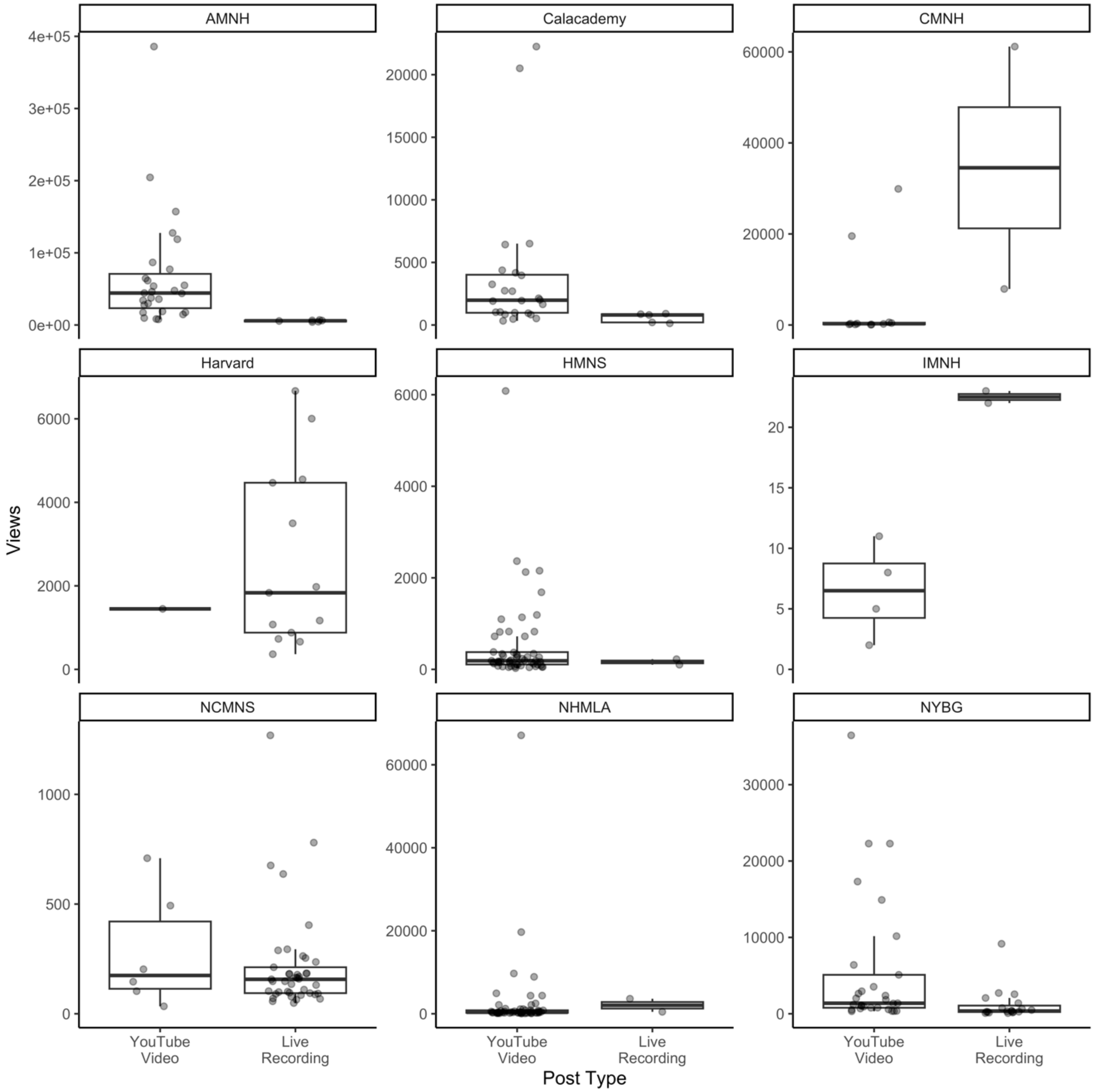


**Supplementary Figure 5.2. Boxplots of the number of video views between YouTube-specific and live recording posts by museum account.** Points represent values from individual videos from each museum jittered horizontally in space. Outliers for the boxplot are suppressed. See Supplementary Table 1.1 for museum acronyms and sample sizes (i.e., number of normal and non-normal videos) for each museum account.

#### 5.3 Number of Likes

Number of likes on a video as of 22 February 2022 was scraped directly from YouTube using YouTube's associated application programming interface (API) for the 2019 dataset. Live recording posts tended to have fewer views than YouTube-specific videos but this was not significantly different when investigating at the entire dataset level ($W = 13{,}502$, $p = 0.01$) and were only significantly different for 3 of the 9 museum accounts that had posted both YouTube-specific videos and live recording posts (Supplementary Table 5.3, Supplementary Figure 5.3). The three museums accounts that were significant all demonstrate that YouTube-specific videos had a higher number of likes than live recording posts.

**Supplementary Table 5.3. Summary of Wilcoxon tests for number of likes between YouTube-specific videos and live recording posts by museum account.** The degrees of freedom indicate the sample size for the live recording and YouTube-specific videos respectively. False discovery rates take into account tests run on video duration, number of video views, and number of likes videos received. **Bold** values are significant.

| **Museum (abbreviation)** | **Statistic (*W*)** | **Degrees of freedom** | ***p* value** | ***q* value** |
| --- | --- | --- | --- | --- |
| American Museum of Natural History (AMNH) | 18.5 | 6, 27 | **0.004** | **0.014** |
| California Academy of Sciences (Calacademy) | 18.0 | 5, 24 | **0.017** | **0.044** |
| Cleveland Museum of Natural History (CMNH) | 20.0 | 2, 11 | 0.092 | 0.177 |
| Harvard Museum of Natural History (Harvard) | 10.0 | 13, 1 | 0.456 | 0.586 |
| Houston Museum of Natural History (HMNS) | 24.5 | 2, 54 | 0.198 | 0.281 |
| Idaho Museum of Natural History (IMNH) | 5.0 | 2, 4 | 0.780 | 0.810 |
| Natural History Museum of Los Angeles County (NHMLA) | 71.0 | 2, 55 | 0.501 | 0.615 |
| New York Botanical Garden (NYBG) | 152.0 | 19, 29 | **0.009** | **0.028** |
| North Carolina Museum of Natural Sciences (NCMNS) | 136.5 | 41, 6 | 0.676 | 0.730 |

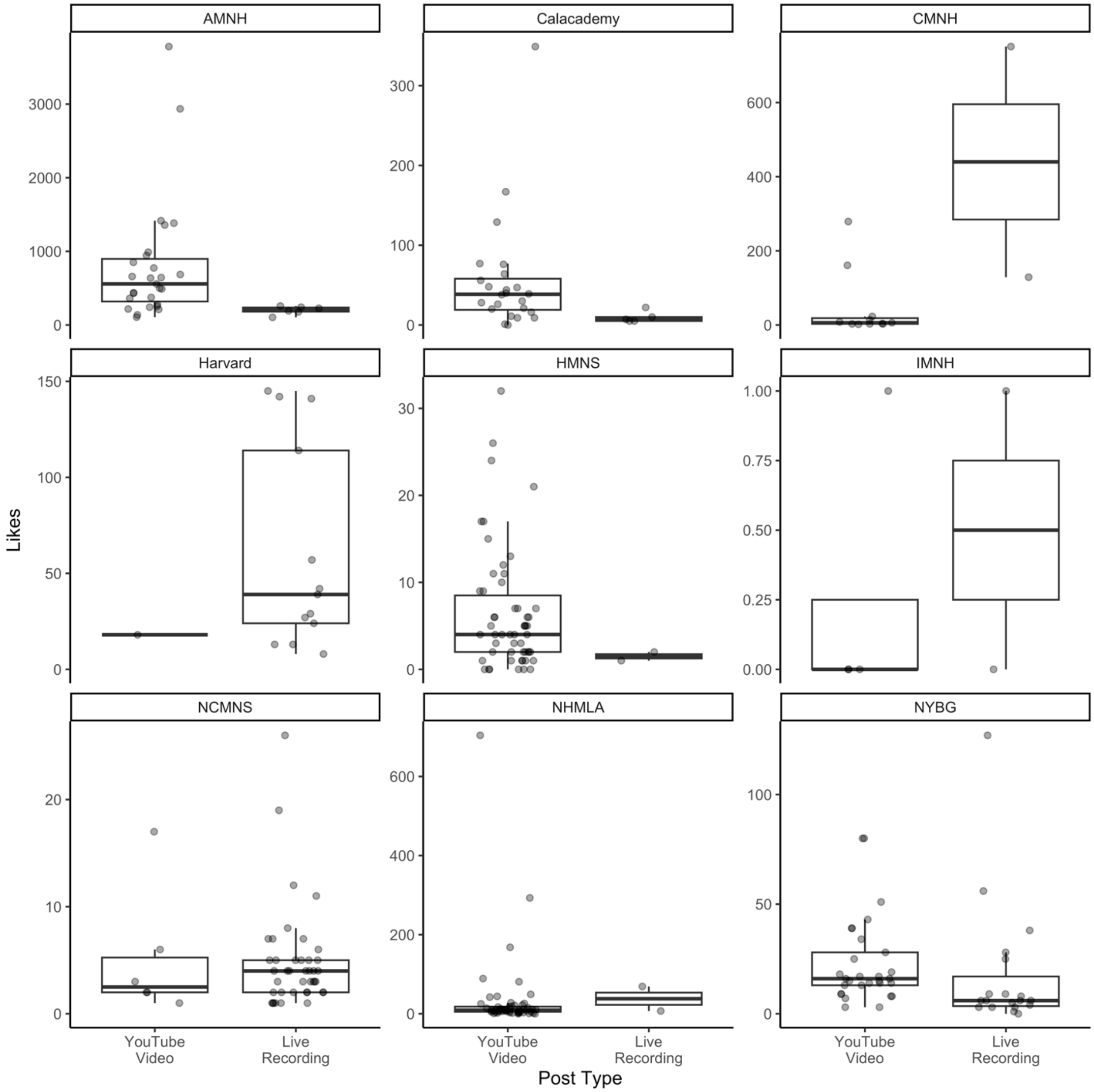


**Supplementary Figure 5.3. Boxplots of the number of video likes between YouTube-specific and live recording posts by museum account.** Points represent values from individual videos from each museum jittered horizontally in space. Outliers for the boxplot are suppressed. See Supplementary Table 1.1 for museum acronyms and sample sizes (i.e., number of YouTube-specific and live recording posts) for each museum account.

### 6 2023 Dataset

#### 6.1 Dataset Summary

The 2023 dataset was manually compiled on 2 August 2023 to determine if trends in both content and production choices between the main dataset (i.e., 2019 dataset) were consistent with recent practices. This dataset consists of fewer videos (134 videos) compared to the 2019 dataset (448 videos) and spans a post time of approximately 3 months (April 2023 through July 2023). Therefore, this dataset is mainly used for comparison and is not the focus of the manuscript. The number of posts with information manually compiled differed depending on the museum (Supplementary Table 6.1).

**Supplementary Table 6.1. Summary information of YouTube museum accounts in the 2023 dataset.** The number of videos posted and between April 2023 through July 2023, along with the number of videos posted in each post type. The number of videos posted and percentage of English YouTube-specific videos in each of the four main topic categories, with first-person language, and with a person on-screen by museum. Num. = Number

| | | **Post Type** | | **Main topic category** | | | | | |
|---|---|---|---|---|---|---|---|---|---|
| **Museum (abbreviation)** | **Num. of posts included in the recent dataset** | **Num. of English YouTube videos** | **Num. of English live recordings (physical presentation, virtual presentation)** | **Num. of videos sharing original research (%)** | **Num. of videos sharing museum/exhibit information (%)** | **Num. of videos sharing specimen/collection information (%)** | **Num. of videos sharing general information (%)** | **Num. of videos with any first-person language (%)** | **Num. of videos with a person on-screen (%)** |
| American Museum of Natural History (AMNH) | 3 | 2 | 1 (1, 0) | 0 (0.0) | 1 (50.0) | 0 (0.0) | 1 (50.0) | 2 (100.0) | 0 (0.0) |
| Arizona Museum of Natural History (AzMNH) | 3 | 3 | 0 (0, 0) | 0 (0.0) | 2 (66.7) | 0 (0.0) | 1 (33.3) | 3 (33.0) | 1 (33.3) |
| Bell Museum (Bell) | 1 | 1 | 0 (0, 0) | 0 (0.0) | 0 (0.0) | 0 (0.0) | 1 (100.0) | 0 (0.0) | 0 (0.0) |
| Bernice P. Bishop Museum (Bishop) | 1 | 0 | 1 (1, 0) | — | — | — | — | — | — |

| | | Post Type | | Main topic category | | | | | |
|---|---|---|---|---|---|---|---|---|---|
| **Museum (abbreviation)** | **Num. of posts included in the recent dataset** | **Num. of English YouTube videos** | **Num. of English live recordings (physical presentation, virtual presentation)** | **Num. of videos sharing original research (%)** | **Num. of videos sharing museum/exhibit information (%)** | **Num. of videos sharing specimen/collection information (%)** | **Num. of videos sharing general information (%)** | **Num. of videos with any first-person language (%)** | **Num. of videos with a person on-screen (%)** |
| Burke Museum of Natural History & Culture (Burke) | 3 | 0 | 3 (1, 2) | — | — | — | — | — | — |
| California Academy of Sciences (Calacademy) | 10 | 6 | 4 (1, 3) | 1 (16.7) | 3 (50.0) | 0 (0.0) | 2 (33.3) | 3 (50.0) | 3 (50.0) |
| Cleveland Museum of Natural History (CMNH) | 1 | 1 | 0 (0, 0) | 0 (0.0) | 1 (100.0) | 0 (0.0) | 0 (0.0) | 1 (100.0) | 1 (100.0) |
| Delaware Museum of Nature & Science (DelMNH) | — | — | — | — | — | — | — | — | — |
| Denver Botanic Gardens (Denver Botanic) | 4 | 3 | 0 (0, 0) | 0 (0.0) | 2 (66.7) | 0 (0.0) | 1 (100.0) | 1 (33.3) | 2 (66.7) |
| Denver Museum of Nature & Science (Denver Museum) | 34 | 22 | 12 (7, 5) | 15 (68.2) | 4 (22.7) | 0 (0.0) | 2 (9.1) | 19 (86.4) | 20 (90.9) |
| Florida Museum of Natural History (Florida Museum) | — | — | — | — | — | — | — | — | — |

| | | **Post Type** | | **Main topic category** | | | | | |
|---|---|---|---|---|---|---|---|---|---|
| **Museum (abbreviation)** | **Num. of posts included in the recent dataset** | **Num. of English YouTube videos** | **Num. of English live recordings (physical presentation, virtual presentation)** | **Num. of videos sharing original research (%)** | **Num. of videos sharing museum/exhibit information (%)** | **Num. of videos sharing specimen/collection information (%)** | **Num. of videos sharing general information (%)** | **Num. of videos with any first-person language (%)** | **Num. of videos with a person on-screen (%)** |
| Harvard Museum of Natural History (Harvard) | 4 | 4 | 0 (0, 0) | 3 (75.0) | 1 (25.0) | 0 (0.0) | 0 (0.0) | 3 (75.0) | 3 (75.0) |
| Houston Museum of Natural History (HMNS) | 5 | 4 | 1 (0, 1) | 0 (0.0) | 1 (25.0) | 0 (0.0) | 3 (75.0) | 4 (100.0) | 2 (50.0) |
| Idaho Museum of Natural History (IMNH) | — | — | — | — | — | — | — | — | — |
| Illinois State Museum (ILState) | — | — | — | — | — | — | — | — | — |
| Kansas University Natural History Museum (KUMHN) | — | — | — | — | — | — | — | — | — |
| La Brea Tar Pits and Museum (La Brea) | 2 | 2 | 0 (0, 0) | 0 (0.0) | 2 (100.0) | 0 (0.0) | 0 (0.0) | 0 (0.0) | 0 (0.0) |
| Natural History Museum of Los Angeles County (NHMLA) | 24 | 19 | 3 (3, 0) | 0 (0.0) | 6 (31.6) | 0 (0.0) | 13 (68.4) | 16 (84.2) | 17 (89.5) |
| New York Botanic Gardens (NYBG) | 14 | 7 | 7 (2, 5) | 1 (14.3) | 6 (85.7) | 0 (0.0) | 0 (0.0) | 3 (42.9) | 2 (28.6) |

| | | **Post Type** | | **Main topic category** | | | | | |
|---|---|---|---|---|---|---|---|---|---|
| **Museum (abbreviation)** | **Num. of posts included in the recent dataset** | **Num. of English YouTube videos** | **Num. of English live recordings (physical presentation, virtual presentation)** | **Num. of videos sharing original research (%)** | **Num. of videos sharing museum/exhibit information (%)** | **Num. of videos sharing specimen/collection information (%)** | **Num. of videos sharing general information (%)** | **Num. of videos with any first-person language (%)** | **Num. of videos with a person on-screen (%)** |
| New York State Museum (NYSM) | 5 | 2 | 3 (3, 0) | 0 (0.0) | 2 (100.0) | 0 (0.0) | 0 (0.0) | 0 (0.0) | 0 (0.0) |
| North Carolina Museum of Natural Sciences (NCMNS) | 5 | 4 | 1 (1, 0) | 0 (0.0) | 4 (100.0) | 0 (0.0) | 0 (0.0) | 3 (75.0) | 2 (50.0) |
| Peggy Notebaert Nature Museum (Nature Museum) | 1 | 1 | 0 (0, 0) | 0 (0.0) | 0 (0.0) | 0 (0.0) | 1 (100.0) | 1 (100.0) | 0 (0.0) |
| Perot Museum of Nature & Science (Perot) | 1 | 1 | 0 (0, 0) | 0 (0.0) | 1 (100.0) | 0 (0.0) | 0 (0.0) | 1 (100.0) | 0 (0.0) |
| Sam Noble Museum (Sam Noble) | — | — | — | — | — | — | — | — | — |
| Smithsonian's National Museum of Natural History (Smithsonian) | 4 | 1 | 3 (0, 3) | 0 (0.0) | 1 (100.0) | 0 (0.0) | 0 (0.0) | 1 (100.0) | 0 (0.0) |
| South Carolina State Museum (SCStateMuseum) | 1 | 1 | 0 (0, 0) | 0 (0.0) | 0 (0.0) | 0 (0.0) | 1 (100.0) | 1 (100.0) | 1 (100.0) |
| The Academy of Natural Sciences (AcadNatSci) | 2 | 0 | 2 (2, 0) | — | — | — | — | — | — |

| | | **Post Type** | | **Main topic category** | | | | | |
|---|---|---|---|---|---|---|---|---|---|
| **Museum (abbreviation)** | **Num. of posts included in the recent dataset** | **Num. of English YouTube videos** | **Num. of English live recordings (physical presentation, virtual presentation)** | **Num. of videos sharing original research (%)** | **Num. of videos sharing museum/exhibit information (%)** | **Num. of videos sharing specimen/collection information (%)** | **Num. of videos sharing general information (%)** | **Num. of videos with any first-person language (%)** | **Num. of videos with a person on-screen (%)** |
| Yale Peabody Museum of Natural History (Peabody) | 6 | 1 | 3 (3, 0) | 0 (0.0) | 0 (0.0) | 3 (100.0) | 0 (0.0) | 1 (33.3) | 3 (66.7) |

#### 6.2 YouTube-Specific Content

Similar to the 2019 dataset, the 2023 dataset also mainly consisted of English posts (3 out of 134 were not in English). Videos considered made for YouTube were still posted more often (66.4%, 87 out of 131 English videos) than videos not initially made for YouTube (44 videos; Figure 1A in main text). Posted content that was not initially made for YouTube again consisted of recordings to physical (56.8%, 25 out of 44 videos) or virtual audiences (43.2%, 19 out of 44 videos) (Figure 1B in main text). Museums again varied in the percentage of their posts that were either YouTube-specific content or live recordings; however, these percentages were not significantly different from the 2019 dataset (Figure 1C in main text, Table 2 in main text, Supplementary Table 6.1).

#### 6.3 Museum Posts and Research and Collections Topics

The main topic categories that museum accounts posted videos in the 2023 dataset followed the same order as when they posted in the 2019 dataset though the percentages of the total dataset did vary: sharing (1) museum or exhibit information (43.7%, 38 posts), (2) general education (29.9%, 26 posts), (3) original research (23.0%, 20 posts), and, least frequently, (4) specimen or collection information (3.4%, 3 posts) (Figure 2A in main text). Across museums this order was still consistent: posting a video that shared museum or exhibit information was still the most common at 53.9% (± standard deviation-SD: ± 40.1%; range: 0.0-100.0%), followed by sharing general information (31.7 ± 38.8%; 0.0-100%), sharing original research (9.2 ± 22.6%; 0.0-75.0%), and lastly by sharing specimen or collection information (5.3 ± 22.9%; 0.0-100.0%) (Figure 2B in main text; Supplementary Table 6.1). In addition to museums only posting videos in the main topic categories of sharing museum or exhibit information and sharing general education, one museum (Yale Peabody Museum of Natural History) posted all three of its videos with a main topic category of sharing specimen or collection information (Supplementary Table 6.1). While these percentages differed from those in the 2019 dataset, there was no significant difference in the trend when accounting for false discovery rates (Table 2 in main text).

There were 21 videos (24.1% of YouTube posts) that featured research. All videos that had a main topic category of sharing original scientific research automatically also had a research section (20 videos). The single remaining video that had a research section was in the main topic category of sharing specimen or collection information. Both sharing museum or exhibit information and sharing general education no longer had videos that also had a research section.

In the 2023 dataset, only 26.4% of videos (23 out of 87) drew attention to collections (either biological or non-biological). This is 1.7 times fewer than in the 2019 dataset. Most commonly, collections were mentioned in sharing museum or exhibit information (10 videos) followed by sharing research (6 videos), sharing general education (4 videos), and lastly, sharing specimen or collection information (3 videos). All the videos that had a main topic category of sharing specimen or collection information drew attention to collections.

#### 6.4 Presentation of Natural History Museum YouTube-Specific Content

Similar to the 2019 dataset, across all post categories and regardless of whether research was mentioned, use of at least one instance of first-person language was common (70.1%). This trend was consistent within museum accounts with museums on average using at least one instance of first-person language 63.9% (± SD: ± 38.0%) of the time in their posts with only three museums never including an instance of first-person language in their posts (Supplementary Table 6.1).

Similar to the 2019 dataset, it was more common for videos to have someone speak on-screen for at least some portion of the video (64.4%). This was not as consistent looking across museums (Person speaks on-screen: mean ± SD: 42.1 ± 38.4%) perhaps because seven museums (compared to only two in the 2019 dataset) did not have anyone appear on-screen. For videos that did not have someone speak on-screen at any point in the video, the 2023 dataset was only slightly more likely to have any off-screen audio narration (54.8%) compared to the 2019 dataset which was more likely to not have any off-screen audio (74.6%) though these differences were not significant (Table 2 in the main text).

Museums included call-to-action items in 50.6% (44 out of 87 videos) of their posts but rarely solicited feedback (1 out of 44 videos).

All of the videos that featured research also included a scientific source (21 out of 21 videos) and a majority of the videos were affiliated with the museum that posted the video (19 out of 21 videos). Eighteen of the videos had a scientist speak on-screen with zero videos having a scientist speak off-screen, and the remaining three videos did not have a scientist speak at all. When the scientist was speaking, they did use at least one instance of first-person language (18 out of 18 videos).

#### 7 Biological and Non-Biological Collections

Biological and non-biological collections were specifically mentioned in 38% and 16%, respectively, of English YouTube-specific videos (Supplementary Figure 7.1A). In the 2023 dataset, these percentages were approximately halved (18.4% and 8.0% for biological and non-biological collections respectively). In both datasets, biological and non-biological collections were mentioned in every main topic category though some categories have higher or lower frequencies (Supplementary Figure 7.1B).

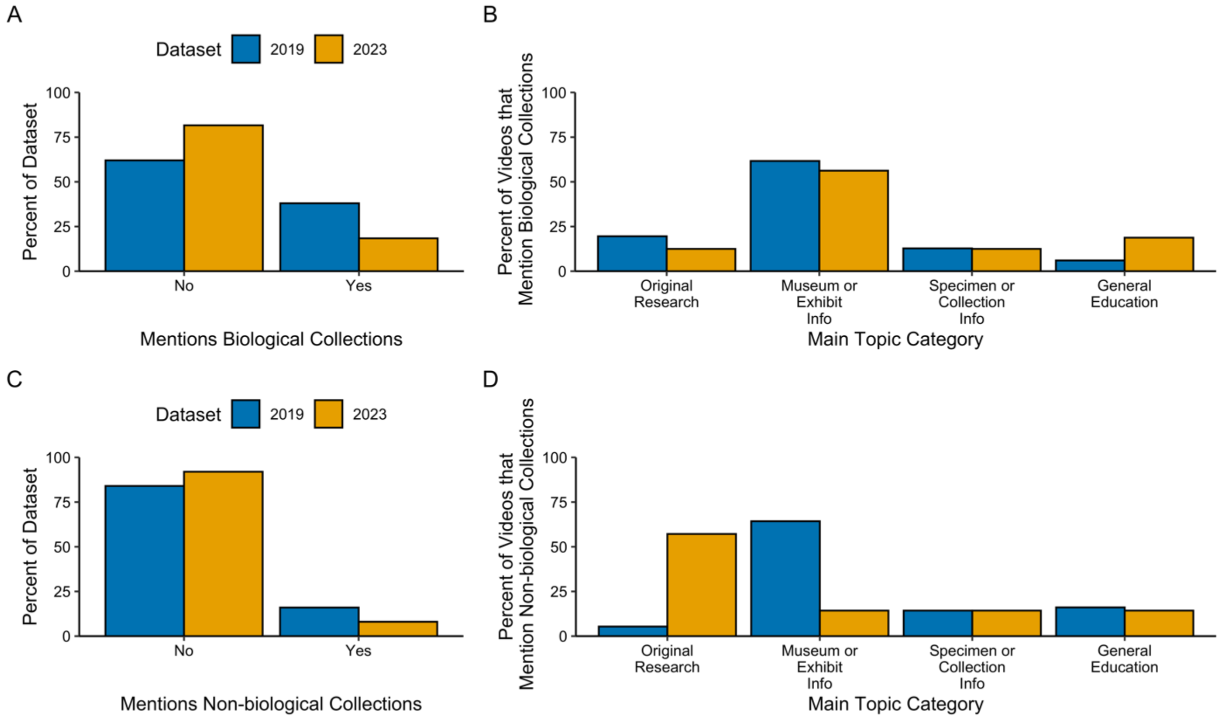

**Supplementary Figure 7.1. Percent of videos that draw attention to biological or non-biological collections by dataset and main topic category.** (A) The overall percentage of normal English videos that draw attention to biological collections at the level of the entire dataset (sample size – videos: 2019 = 350, 2023: 87). (B) Percentage of videos that draw attention to biological collections in each main topic category at the level of the entire dataset (sample size - videos: 2019 = 133, 2023 = 16). (C) The overall percentage of normal English videos that draw attention to non-biological collections at the level of the entire dataset (sample size – videos: 2019 = 350, 2023: 87). (D) Percentage of videos that draw attention to non-biological collections in each main topic category at the level of the entire dataset (sample size - videos: 2019 = 56, 2023 = 7).
